# Uncovering distinct protein conformations using coevolutionary information and AlphaFold

**DOI:** 10.1101/2025.10.08.681198

**Authors:** Yongkai Chen, Samuel W.K. Wong, S. C. Kou

## Abstract

Protein structure prediction has been transformed by AlphaFold, yet a key challenge remains: characterizing the multiple conformations adopted by proteins that can switch between different folds, without knowledge of their potential binding partners. Existing methods rely on sampling the multiple sequence alignment (MSA), either through random sampling or clustering, but these methods are statistically inefficient and do not explicitly utilize coevolutionary information during MSA sampling. We introduce an iterative sampling framework that systematically explores the MSA space using residue-specific frequencies and coevolutionary patterns inferred via Markov random fields. We further develop tools to identify a protein’s variable region and extract representative structures, yielding a compact, high-quality ensemble with good coverage of distinct conformations. On a benchmark set of fold-switching proteins, our method outperforms existing ones by substantially improving the diversity of the sampled structures. Overall, this work significantly advances our ability to characterize the conformational landscape of proteins.

## Introduction

It is fair to say that AlphaFold2 (*1*) has revolutionized the task of protein structure prediction from sequences. While AlphaFold2 has received incremental updates since its inception (*2, 3*) and continues to be ubiquitous for structure prediction of single protein sequences, it is still challenging to adequately characterize distinct conformational states for a given protein. Many proteins, such as fold-switching or metamorphic proteins, are dynamic and change in response to their environment, multimerization, and/or binding partners (*4, 5*). In this regard, AlphaFold3 (*6*) enhanced the prediction of specific protein complexes and binding-induced conformational changes. On the other hand, accurately predicting the possible conformational switches for a protein, without knowledge of its potential binding partners, represents a substantially more difficult problem that has profound applications (*7*). For example, traditional drug design focuses on a single, static target protein. However, if the target protein can exist in multiple conformations, a drug that targets one state may be ineffective or even harmful if it (unintentionally) drives the protein toward a pathogenic state (*8*). Ideally, we would like to characterize the conformational landscape accessible to a given protein across its unbound and bound conditions. However, each entry in the Protein Data Bank [PDB, (*9*)] only provides a static experimentally-determined structure for most sequences; for proteins whose structures have been solved multiple times, the set of static experimental structures often confirms the presence of conformational variability (*10*). Laboratory techniques to capture protein conformational changes remain limited, and this limitation extends to AlphaFold, which is trained under the one-sequence-to-one-structure paradigm, assuming the protein will stabilize at its lowest-energy conformation in accordance with the energy landscape theory (*11–13*). When predicting the structures of fold-switching proteins, which have at least two distinct yet stable structures, AlphaFold2 by default can only predict one of the structures (*14*).

This limitation has inspired studies into how AlphaFold2 can be enhanced to provide predictions that capture all possible foldings for a protein sequence. These efforts largely revolve around modifying the input to AlphaFold2, most notably in the form of the multiple sequence alignment (MSA). An MSA consists of a list of protein sequences that are evolutionarily or structurally related to the target sequence, usually retrieved by querying large databases of known protein sequences (*15–17*). With a well-constructed MSA, AlphaFold2 is generally regarded to be more accurate than protein language model-based folding algorithms that operate on the input sequence only, e.g., the ESM family (*18*) and OmegaFold (*19*). Existing approaches to diversify AlphaFold2’s predictions have focused on sampling the full MSA (or using multiple shallow MSAs) to generate structures from a larger conformational space. These include *random sampling* (*20–22*), which draw fixed-size random subsets from the full MSA, and *AF-Cluster* (*23*), where sequences similar to each other are grouped into clusters that are used as input MSAs to AlphaFold2.

However, these approaches have two main limitations. First, from a statistical perspective, they do not explore the space of MSA subsets in an effective or unbiased manner. The ideal case for fully leveraging prediction diversity via MSA sampling would be to exhaustively enumerate all possible subsets, ranging from subsets consisting of diverse sequences to subsets with highly homogeneous sequences, along with intermediate combinations. Clustering methods are inherently biased as they only sample those subsets with highly homogeneous sequences. In contrast, random sampling, though theoretically unbiased, suffers from low sampling efficiency and does not use any sequence information within the MSA. Second, AF-Cluster treats each residue in the sequence independently when constructing MSA subsets: it compares the amino acid types for each residue marginally and disregards the interaction between different residues in the selection of MSA subsets. Coevolutionary information, which refers to the statistical dependence between two residue positions (potentially far apart), is believed to play an important role in how AlphaFold makes predictions (*24*). Emerging evidence also suggests that the prediction of multiple conformations through MSA sampling benefits from selecting subsets that contain distinct coevolutionary information patterns (*23*). However, existing approaches, including AF-Cluster and random sampling, do not explicitly incorporate coevolutionary information in MSA sampling.

To address these limitations, we propose an iterative sampling method, SMICE (**S**ampling **M**SA **I**teratively with **C**o**E**volution information), which formally embeds MSA sampling into generative probabilistic models and incorporates the coevolutionary information into the MSA sampling criterion. The method begins with sequentially sampling sequences based on each residue’s marginal frequencies using Bayesian configurations that ensures high diversity across all sampled subsets. Subsequently, the sampled MSA subsets are used in AlphaFold2 to generate an initial set of structure predictions. Next, to increase the diversity of MSA subsets’ coevolutionary information, we extract the structurally distinct predictions and profile the coevolutionary information from each corresponding MSA subset using a Markov random field (MRF) model (*25*). This approach efficiently captures variations in the coevolutionary information that contributes to the structural diversity of the predictions. The fitted MRF models then guide the construction of a new round of MSA subsets with increasingly diverse coevolutionary information. These newly constructed MSA subsets are subsequently used in AlphaFold2 to predict structures. This process is iterated to generate a sufficiently large number of predictions.

During this iterative process, SMICE usually generates a large number of structure predictions (typically more than 1,000) to ensure thorough exploration of the conformational space. Extracting a compact set of representative structures is thus a critical next step for efficient downstream analysis. While previous studies have focused on using AlphaFold’s confidence metrics (e.g., AlphaFold2’s built-in learned predicted local distance difference test, pLDDT) to rank predictions (*23, 26*), we found this approach problematic for fold-switching proteins due to a tendency for the confidence metrics to be overestimated (Supplementary Material, Figure S4, S5). In contrast, we developed an integrated procedure for extracting representative structures that combines confidence metrics with structural clustering. This combined strategy efficiently reduces the number of final structures for examination from over 1,000 to fewer than 30 in most cases, while maintaining high-quality structures that ensure good coverage of the distinct conformational states.

Our results demonstrate the advantages of enhancing the diversity of MSA subsets using both amino acid frequencies and coevolutionary information. For example, SMICE successfully captures both conformational states of the fold-switching protein RelB, whereas AF-Cluster and random sampling are unable to predict the fold-switching state (as will be discussed shortly in the Results section). The performance of SMICE is demonstrated on a benchmark set of fold-switching proteins from (*26*), where each protein is known to have two distinct conformations. In this benchmark dataset, SMICE consistently outperforms other methods in predicting both conformations while maintaining high prediction fidelity across most cases. Overall, our method provides a reliable solution for predicting multiple conformations of proteins capable of adopting distinct folds. These insights could lead to a deeper subsequent understanding of conformational changes and the critical role of fold-switching in protein function and evolution.

## Results

### Framework of SMICE

Our method consists of two steps: the sampling step (Fig. 1a) and the representative extraction step (Fig. 1b). In the sampling step, a statistical sequential sampling procedure is first applied to the full MSA to produce MSA subsets with diverse marginal statistics (amino acid proportions per residue), driven by a Bayesian framework. When sampling each different MSA subset, the sequence sampling probability is computed under a Bayesian prior distribution of amino acid proportions. Varying the Bayesian prior distribution enables a broader exploration of the conformational space by sampling MSA subsets with distinct conservation patterns in both the locations of conserved residues and the types of enriched amino acids. Next, the method leverages coevolutionary information that would not have been captured in the marginal statistics used by sequential sampling. This begins with selecting those MSA subsets that predict the most structurally diverse conformations. Then, to utilize and enhance the differences in the coevolutionary information of these representative MSA subsets, a Markov Random Field (MRF) model (*25*) is fitted to each of the MSA subsets. We then rank sequences from the full MSA by their probability ratios under these competing MRF models. By selecting sequences that strongly favor one model over another, we construct new MSA subsets enriched with specific coevolutionary information. This process is iterated to ensure thorough exploration of the conformational space. The predicted structures from both the sequential sampling and the enhanced (coevolution-aware) sampling are combined as the sampling result of SMICE.

**Figure 1.**
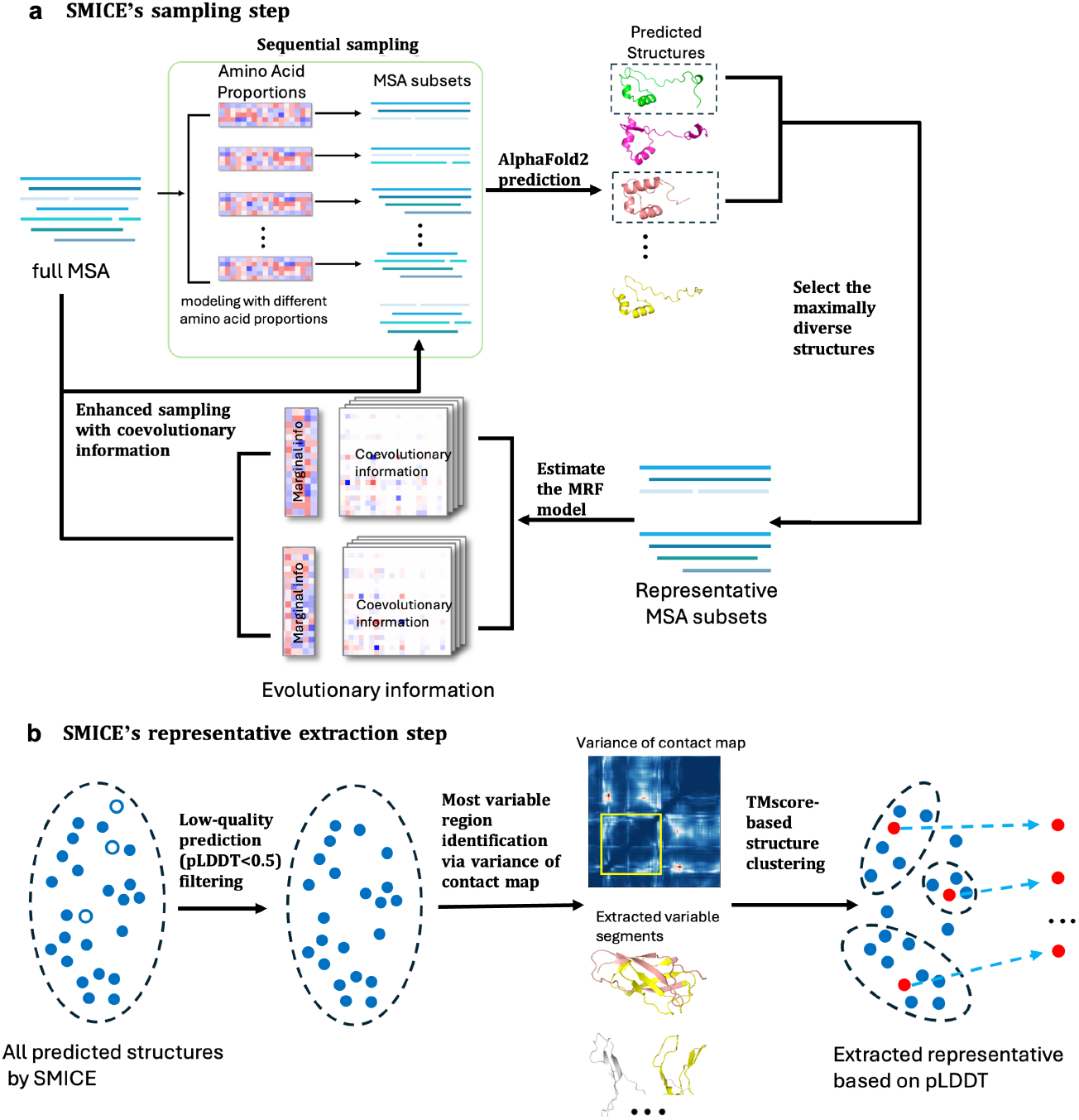
SMICE workflow. **a**, Flowchart of SMICE’s sampling step. MSA subsets are drawn from the full MSA using sequential sampling, which are initialized by modeling MSA subsets with different amino acid proportions. Then, structure predictions are made on the MSA subsets with AlphaFold2. Representative MSA subsets are selected by maximizing the diversity of their corresponding structures. For each representative MSA subset, we estimate its coevolutionary information using a Markov random field (MRF) model. The next round of MSA subsets is constructed via sampling, which utilizes the differences in coevolutionary information embedded within the representative MSA subsets. **b**, Flowchart of SMICE’s representative extraction step. First, low-quality predictions are filtered by a pLDDT threshold of 0.5. Next, the variance of the contact map is calculated across the remaining structures to identify the variable region. Predicted structures are then segmented according to the variable region, and clustered by TMscores, calculated on the variable region plus its neighboring segments, to identify groups and outliers. The structure with the highest pLDDT score within each cluster is chosen as its representative.

The second step of SMICE is extracting a compact set of representative structures from all the predictions. The procedure (Fig. 1b) is designed as follows: First, low-quality predictions are filtered out based on the pLDDT scores. An analysis of filtering metrics and threshold choices is provided in the Supplementary Material. Then, the variance of the residue contact map is calculated across the remaining structures, and the variable region of the protein is identified (the definition of the variable region and the algorithm are provided in the Method section). Next, we cluster the high-quality structures based on their structural similarity in the variable region. After identifying the clusters and excluding the outliers, the structure with the highest pLDDT score within each cluster forms the final set of representative structures. A detailed description of SMICE is provided in the Method section.

### Performance of SMICE’s sampling step on RelB

We consider the example of predicting the conformational states of the transcription factor RelB to illustrate the performance of SMICE’s sampling step (Fig. 1a). RelB adopts two distinct structural states, each corresponding to a different dimerization mode (*27*): (1) heterodimerization with other NF-*κ*B subunits (ground state, Fig. 2b, PDB ID, 3jv6A) or (2) homodimerization (transiently stable but not observed *in vivo*, Fig. 2b, PDB ID, 1zk9A).

**Figure 2.**
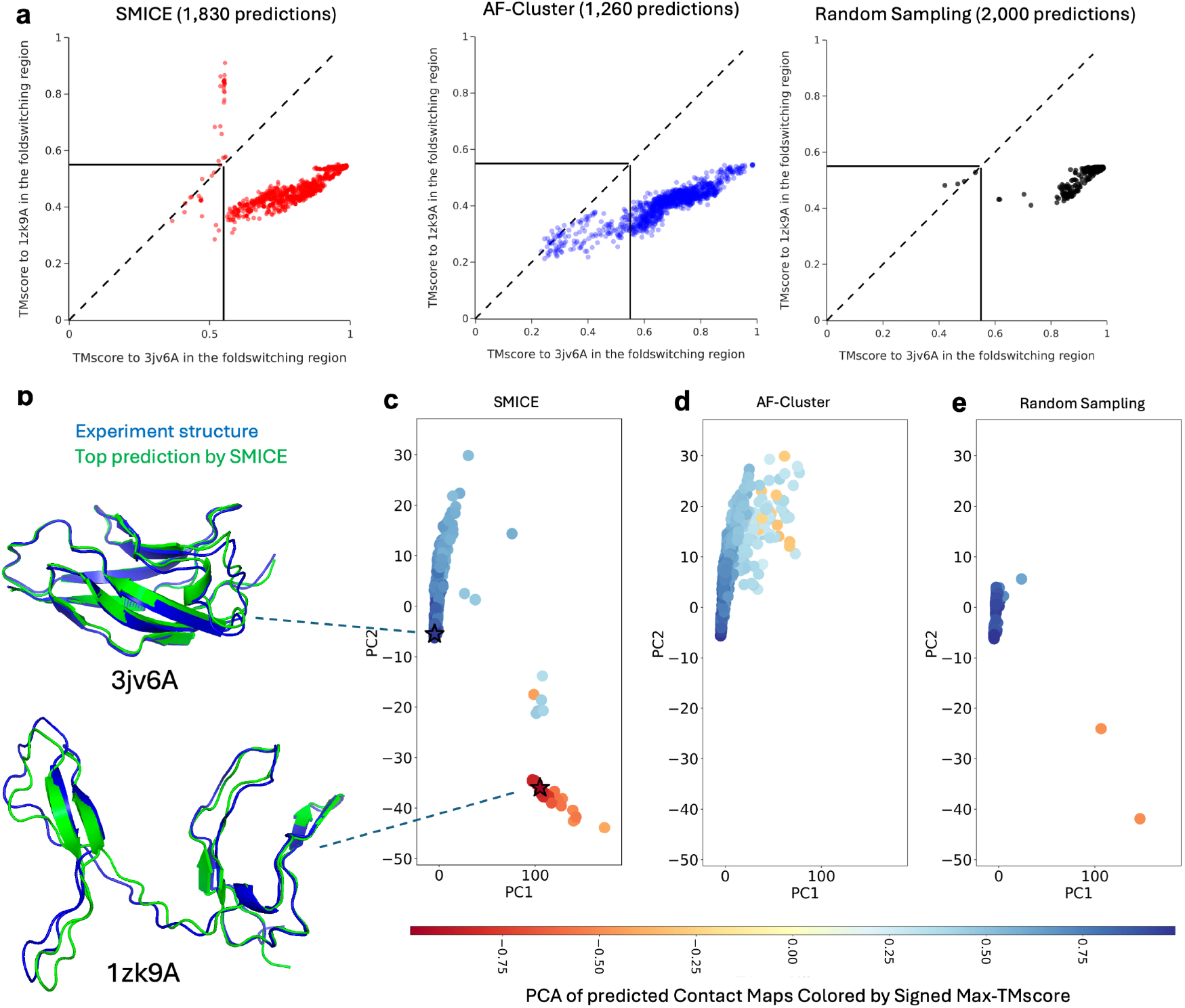
SMICE’s sampling captures both known conformations of the fold-switching protein RelB. **a**, TMscores of AlphaFold2 predictions using MSA subsets generated by SMICE (red), AF-Cluster (blue), and random sampling (black). The TMscore is calculated on the fsr plus its neighboring segments. Black vertical and horizontal lines indicate TMscores between RelB’s two conformations, 3jv6A and 1zk9A. **b**, Crystal structures of RelB in its ground state (PDB: 3jv6A) and fold-switching state (PDB: 1zk9A) alongside their top SMICE-predicted structures. **c-e**, Principal component analysis (PCA) of predicted contact maps from SMICE (**c**), AF-Cluster (**d**), and random sampling (**e**). Each point represents the first two principal components of each predicted structure’s contact map, colored by the signed max-TM score (sign[TMscore to 3jv6A minus TMscore to 1zk9A] × max of TMscores to 3jv6A and 1zk9A). Star markers denote the two conformational states 3jv6A (blue) and 1zk9A (red).

To compare the efficacy of different methods, we ran SMICE, AF-Cluster, and random sampling on the full MSA of RelB (see Supplementary Material for implementation details) to predict its conformational states. To assess prediction accuracy, we computed TMscore, a widely used metric for evaluating structural similarity between two structures, for a combined region that includes the fold-switching region (fsr), as annotated by (*28*), *and* its neighboring left and right segments, each with a length equal to half that of the fsr. As noted by (*26*), whole-protein TMscores tend to overestimate prediction accuracy; furthermore, we observed that TMscores calculated solely on the fsr often fail to distinguish conformations resulting from hinge motions (*29*), where the global fold remains largely unchanged but relative orientations between regions shift. Therefore, we based the TMscore calculation on the fsr plus its neighboring segments.

Notably, only SMICE successfully generated predictions capturing the fold-switching state 1zk9A (Fig. 2a). The top predictions for the state 3jv6A (TMscore: 0.989) and 1zk9A (TMscore: 0.910) by SMICE are visualized in Fig. 2b. Corresponding results, which instead use the whole-structure TMscore and whole-structure RMSD as comparison metrics, are provided in Figure S8 and support the same conclusion.

We were interested in the distribution of predicted structures from the three competing methods. We projected the contact maps of all predictions from the three methods into the principal component analysis (PCA) space (Fig. 2c-e). SMICE predictions are clearly separable into two clusters, each corresponding to a conformational state of RelB. In contrast, AF-Cluster and random sampling did not generate structures close to the fold-switching state 1zk9A. AF-Cluster also produced a significant number of predictions with low TMscores to either of the known states, whereas the predictions from random sampling lacked structural diversity.

### Extracting Representative Structures from SMICE Predictions for RelB

We next extracted representative structures for RelB. This step (Fig. 1b) identified eighteen clusters, each with a representative structure. As visualized by the PCA embedding of these representative structures’ contact maps (Fig. 3a), the representative structures successfully captured the conformational landscape sampled by the predictions. Moreover, clustering revealed an imbalanced prediction set, with a dominant cluster of over 1,500 structures corresponding to RelB’s ground state and several smaller clusters of approximately 20 structures each. Extracting representative structures helped mitigate this imbalance. Notably, the fifth representative structure identified by SMICE corresponds to the known fold-switching state 1zk9A of RelB.

**Figure 3.**
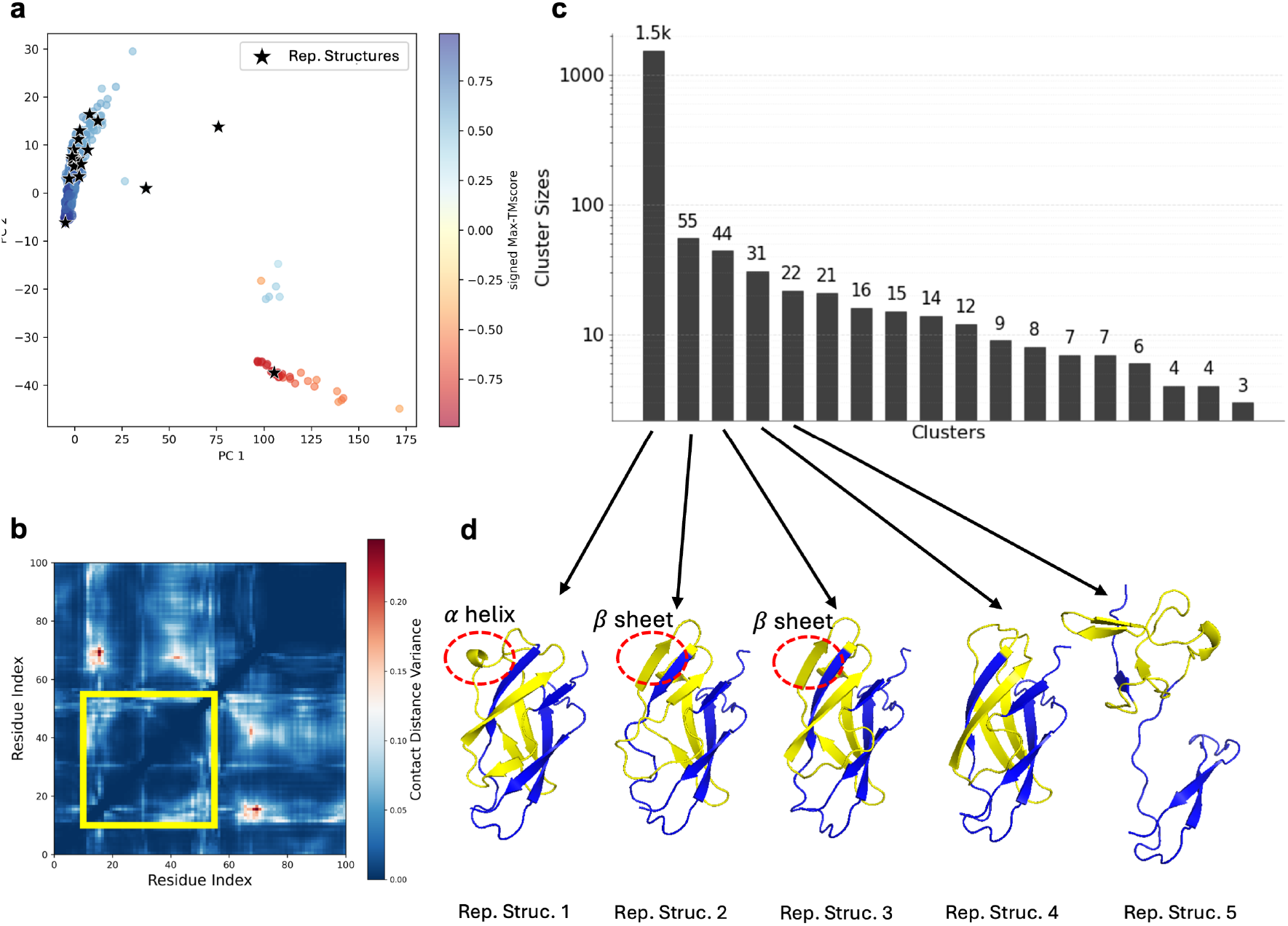
**a**, PCA of predicted contact maps from SMICE on RelB with the representative structures being marked by the stars. **b**, The heatmap of the variance of the predicted contact maps. The yellow rectangle highlights our identified variable region. **c**, The barplot of the cluster sizes, with the y-axis on a logarithmic scale. **d**, The top five representative structures. The variable region is colored yellow.

Comparison of the top five representative structures using our identified variable region (Fig. 3b) reveals three key findings. First, the primary source of variance is the contact distances between the variable region and the rest of the protein, which reflects the hinge motion of the variable region from the ground state 3jv6A to the fold-switching state 1zk9A. Second, our identified variable region well captured the region with the conformational changes. Third, the overall folds of the first four representative structures are similar but exhibit slight secondary or tertiary structural changes within the variable region. For example, an *α*-helix in representative structure 1 changes to a *β*-sheet in representative structures 2 and 3, which are potential conformations that have not been reported in the PDB.

### Experiments on a Fold-Switching Protein Benchmark

We next conducted a systematic examination of the performance of SMICE on a benchmark set of 92 fold-switching proteins provided in (*26*).

We begin by comparing the proportion of successful predictions generated by each method. The prediction set of SMICE for each fold-switching protein consists of all predictions from SMICE’s sampling step, as shown in Fig. 1a. For each pair of conformations of a given fold-switching protein, we use the Fold1 and Fold2 designation by (*26*), where the conformation with higher TMscores for at least 3 out of 5 of their default AlphaFold2 predictions is designated Fold1 and the other conformation is denoted as Fold2, which is more challenging to predict. First, taking each set of predictions generated for a given fold-switching protein, we check if the structures (1) predict Fold1, predict Fold2, or (3) fail to predict either fold. To measure these, we compare the TMscores of both folds against a predefined threshold: a prediction is considered to successfully predict Fold1 if TMscore1 (TMscore to Fold1) exceeds both TMscore2 (TMscore to Fold2) and the threshold, to successfully predict Fold2 if TMscore2 exceeds both TMscore1 and the threshold, and as a failure if neither TMscore1 nor TMscore2 reaches the threshold. Here, the TMscore is calculated on the fsr plus its neighboring segments. By varying the threshold, we calculate the proportions of predictions of these three cases across different levels of prediction confidence for each competing method. Intuitively, only those prediction sets with high proportions for both predicting Fold1 and Fold2 can be considered as successful. Therefore, we use the minimum of the two prediction proportions as a performance metric, representing the ability to successfully predict both folds simultaneously.

In Fig. 4a-c, we summarize the results across all fold-switching proteins with the median of the successful prediction proportions for Fold1, the median of the successful prediction proportions for Fold2, and the median of the minimum proportions for predicting both folds. The plotted curves represent the proportions over the fold-switching proteins, as a function of the TMscore threshold. It is seen that SMICE outperforms AF-Cluster and random sampling, achieving the highest prediction proportions for both folds and the highest minimum success rates across all the thresholds over 0.3. An interesting finding is that random sampling performs comparably to SMICE in predicting Fold1. This may be due to the random sampling’s tendency to generate MSA subsets that share similar information with the full MSAs, leading to their predictions being closer to the more easily predictable Fold1 conformation.

**Figure 4.**
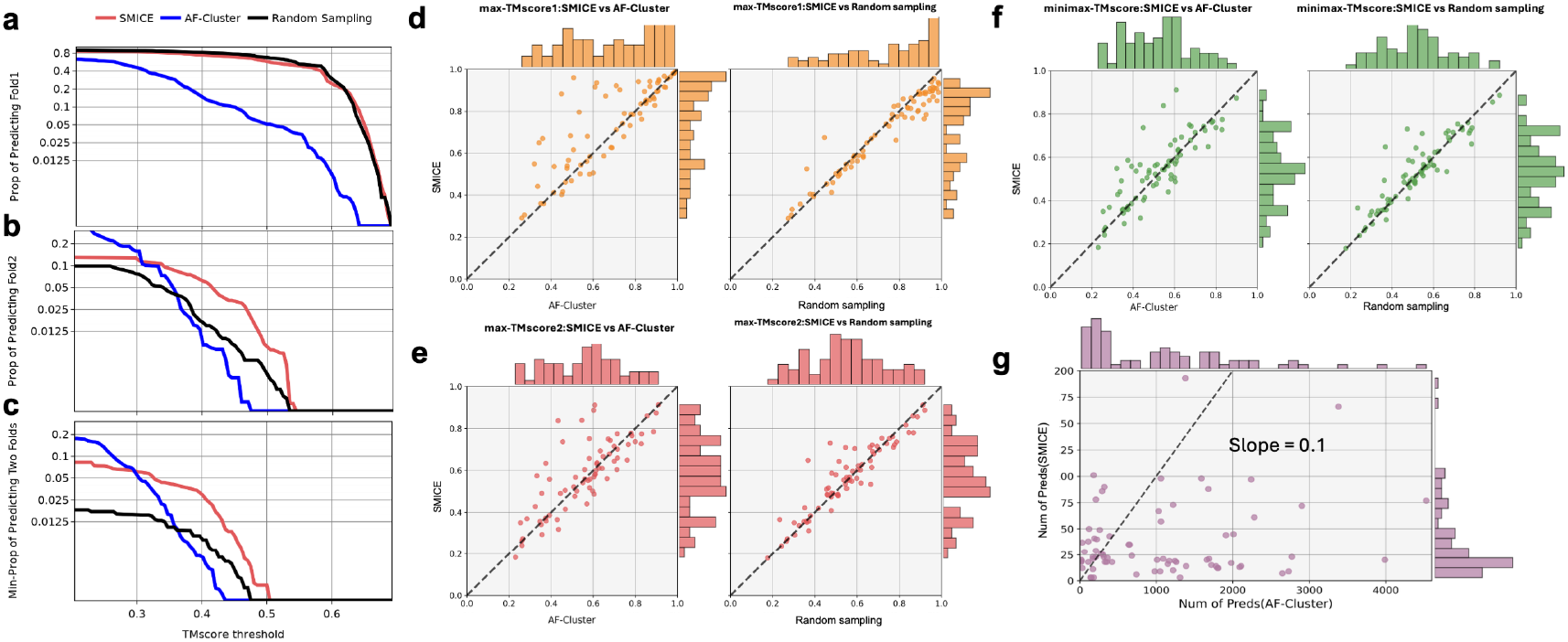
**a-c**, Comparison results of predictions from SMICE’s sampling step against AF-Cluster and Random Sampling on the proportions of predicting Fold1, the proportions of predicting Fold2, and the minimum proportions of predicting both folds. For varying TMscore thresholds, we compute the median of the proportions across all fold-switching proteins. The TMscore is calculated on the fsr plus its neighboring segments. Different methods are colored differently. **d-f**, Comparison of the best prediction result for each method on each fold-switching protein. The top predictions of SMICE are from the final selected representative structures. The rows display the three evaluation metrics: **d**, max-TMscore1; **e**, max-TMscore2; and **f**, minimax-TMscore. In each row, the left panel shows the comparison between SMICE and AF-Cluster, while the right panel shows the comparison with random sampling. Each point represents one fold-switching protein. **g**, Comparison of the number of representative structures of SMICE against the number of predictions by AF-Cluster.

We now compare the best prediction result for each method on each fold-switching protein. For each fold-switching protein, we compute three metrics on its prediction sets: (1) max-TMscore1, i.e., the maximum TMscore to Fold1; (2) max-TMscore2, i.e., the maximum TMscore to Fold2; minimax-TMscore, i.e., the minimum of max-TMscore1 and max-TMscore2, reflecting the worst-case prediction gap for either conformation. A high minimax-TMscore means that both folds are accurately predicted. The TMscore is calculated on the fsr plus its neighboring segments. For SMICE, we used the final predictions after its representative extraction step (Fig. 1b). The best prediction results from SMICE’s sampling step are provided in Fig. S9, which shows even higher prediction accuracy. The results comparing the top predictions of SMICE against the top predictions from AF-Cluster and random sampling are shown in Fig. 4d-f. We found that SMICE outperformed AF-Cluster in capturing both conformations in most cases. Random sampling performed comparably to SMICE for predicting Fold1 (structures that are easier to predict using AlphaFold2 without sampling), while it was outperformed by SMICE in predicting Fold2.

In Fig. 4g, we compare the number of representative structures identified by SMICE against the number of predictions generated by AF-Cluster for each fold-switching protein. We find that the number of structures finally selected by SMICE is typically below 30, with two outliers at approximately 200. In contrast, the number of predictions from AF-Cluster is generally 10 times larger than that of SMICE, with extreme cases yielding over 4,000 predictions per protein. In summary, SMICE outperformed the other methods in terms of both overall prediction quality and the accuracy of the top prediction.

## Discussion

Our work addresses a critical limitation of AlphaFold in predicting the multiple conformations of proteins, without knowledge of potential binding partners. By reformulating MSA sampling as a probabilistic, coevolutionary information-aware procedure, we avoid the tendency of random sampling to choose uninformative MSA inputs and the tendency of clustering to choose only highly homogeneous sequences. For the thioesterase protein SP 1851, we found that MSA subsets with different levels of sequence diversity also exhibit distinct conformational prediction preferences (Figure S1). This direct relationship demonstrates that sampling MSA subsets with multiple levels of sequence diversity (via Bayesian prior distributions) ensures broader coverage of the conformational space. Explicitly leveraging coevolutionary information (by modeling residue dependence) further increases the diversity of coevolutionary information in the MSA subsets. In the analysis of diheme cytochrome c peroxidase from Paracoccus pantotrophus, we found that sequential sampling alone, which only uses the marginal statistics of the MSA, did not capture the fold-switching state (Figure S2). Similarly, AF-Cluster and random sampling, which do not explicitly incorporate coevolutionary information during MSA sampling, could not predict the fold-switching state (Figure S3). In contrast, the full sampling procedure of SMICE, which incorporates residue dependencies and diverse coevolutionary information in the sampled MSA subsets, did successfully predict both conformations.

More ablation studies on the role of coevolutionary information are provided in the Supplementary Material. Our findings demonstrate that MSAs may embed distinct evolutionary and coevolutionary signals, and effectively using this information can significantly improve the diversity and accuracy of predicted conformations.

Extracting representative structures from a large ensemble of predicted structures is essential for characterizing the conformational states of proteins that can switch folds. Since these proteins typically exhibit only a few stable states, identifying these conformations could avoid the need to examine every predicted structure in downstream analyses. SMICE provides a systematic framework by identifying variable regions in the predicted structures, leveraging clustering and AlphaFold’s confidence metrics, to extract representative structures. In benchmark studies, this approach effectively characterized the conformational landscape of most proteins using fewer than 30 representative structures, reduced from an ensemble of over 1,000 predictions.

Fold-switching proteins are likely widespread, but identifying them among the millions of known proteins is challenging since experimental validation is infeasible at such a scale. The presence of heterogeneous coevolutionary information within an MSA, as identified by SMICE, could serve as a computational marker for identifying fold-switching proteins. After detecting strong but diverse coevolutionary information, SMICE could generate a compact set of distinct structures to use for concrete, testable hypotheses. This integrated computational approach could greatly accelerate the discovery of novel fold-switching proteins by focusing costly experimental efforts on high-probability targets. Furthermore, understanding the evolutionary signals that enable fold-switching would facilitate the discovery of de novo protein switches, which in turn could find broad applications in the development of biosensors or targeted drug delivery systems.

## Author contributions

S.K., S.W. and Y.C. designed research; Y.C., S.W. and S.K. performed research; Y.C. implemented the algorithms and analyzed data; and Y.C., S.W. and S.K. wrote the paper.

## Data and materials availability

Data and code corresponding to SMICE and analysis presented here are publicly available at GitHub (https://github.com/StatCYK/SMICE).

## Use of ChatGPT

During the preparation of this manuscript, the authors used ChatGPT (https://openai.com/) solely for improving the language, readability, and correcting the grammar.

## Supplementary Materials

### Method

The first three subsections describe the sampling step of SMICE. The two subsequent subsections describe the representative extraction step of SMICE.

#### Sequential Sampling on the Full MSA

In this section, we develop a sequential sampling procedure to create MSA subsets with diverse marginal statistics, i.e., the amino acid frequencies of each residue. Let ℳ = {**Y**_1_, …, **Y**_*n*_} denote the set of sequences in the full MSA for the target sequence, where **Y**_*i*_, a *L* × 22 binary matrix, is the one-hot encoding of the *i*^*th*^ sequence. The 22 categories include the 20 standard amino acids, one for unknown amino acid types, and one for gaps in the alignment. Suppose we start with a small MSA subset 𝒜 ⊂ ℳ. To facilitate preservation of the evolutionary information in 𝒜, our sequential sampling procedure randomly draws a new (i.e., unsampled in the current MSA subset) sequence 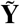 with an acceptance probability proportional to its similarity to the current MSA subset 𝒜. To achieve this, we design the acceptance probability of 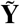 based on how its inclusion changes the amino acid proportions at all residues of 𝒜.

Let ***p***_*l*_ be the length-22 vector of probabilities for the different amino acid types at the *l*^*th*^ residue position, *l* = 1, …, *L*. For an initialization of 𝒜 starting from ∅, we estimate 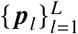 using a Bayesian approach with a prior distribution specified by a *L* × 22 matrix **Π**, where the *l*^*th*^ row of **Π** is the prior vector for the *l*^*th*^ residue position. Specifically, we model that ***p***_*l*_ independently follows a Dirichlet distribution,

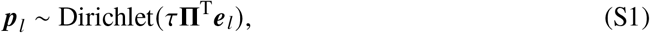

where *τ* > 0 controls the strength of the prior and ***e***_*l*_ is the *l*^*th*^ unit vector so that **Π**^T^***e***_*l*_ gives the *l*^*th*^ row of **Π**.

Given the current MSA subset 𝒜 and the prior distribution of 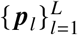 in Eq.(S1), the maximum a posteriori (MAP) estimate of ***p***_*l*_ is

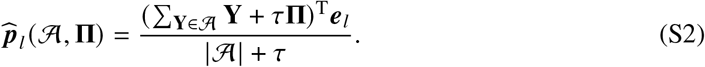

Notice that in the early stages of sequential sampling, the MAP estimate is dominated by **Π**. By assigning high sampling probability to sequences that make small changes to the MAP estimates, sequences that closely match **Π** will be more favored. This allows us to generate MSA subsets covering different local regions in the sequence space by varying the choices of **Π**.

To decide whether to accept a candidate sequence 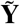, we first calculate the *L*_1_ norm of the MAP estimate’s change by including 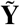 vs. not including it,

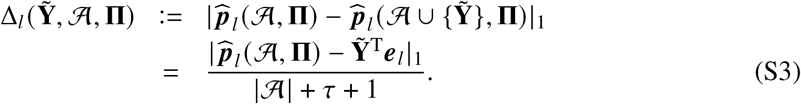

To measure the overall change of the MAP estimate of 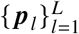, we consider two sources of heterogeneity across all residues: (1) different residues have different levels of evolutionary conservation, and (2) residues with many gaps may provide less reliable information. To address them, we rescale the changes for each residue through

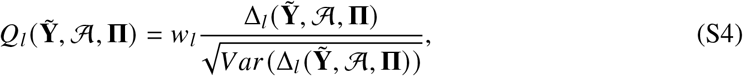

where *V ar* 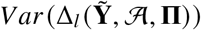 is computed by assuming the marginal distribution of the *l*^*th*^ row of 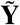 is the one-trial multinomial distribution with probability vector 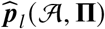, and *w*_*l*_ is the proportion of gaps in the *l*^*th*^ position for the full MSA. By assuming the marginal distribution of the *l*^*th*^ row of 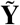 is the one-trial multinomial distribution with probability vector as 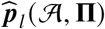, we have

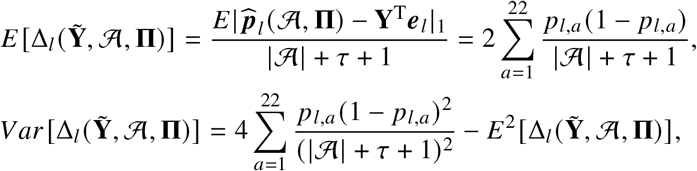

where *p*_*l, a*_ is the *a*^*th*^ entry of 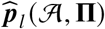.

We now calculate the acceptance probability of the randomly drawn candidate sequence 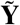 given the current MSA subset 𝒜 through

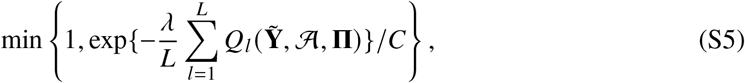

where *C* is a tuning parameter controlling the overall sample size, and *λ* ≥ 0 controls the homogeneity level in the sampled MSA subsets. When *λ* is larger, only sequences resulting in small changes to the MAP estimate are likely to be accepted. This leads to highly homogeneous subsets where sampled sequences are tightly clustered. In contrast, a small *λ* allows for greater diversity in the sampled MSA subsets. In the limiting case of *λ* = 0, sequential sampling reduces to uniform random sampling. For broader exploration, we recommend sampling with multiple *λ* values.

After we query all the sequences, or the size of the MSA subset 𝒜 reaches the preset maximum size, we use 𝒜 as the input MSA for AlphaFold2. By repeating this procedure multiple times with varying choices of the hyperparameters **Π** and *λ*, we induce a set of MSA subsets and their corresponding AlphaFold2 predictions.

#### Selection of Representative MSA Subsets

From the sequentially sampled MSA subsets, we then select the representative MSA subsets based on their structure predictions. Given the AlphaFold2 predictions obtained in the previous step, we expect that notably distinct structures also exhibit distinct evolutionary information within their corresponding MSA subsets. Thus, selecting representative structures and their MSA subsets could provide valuable information for further generation of new MSA subsets that lead to more diverse conformations.

Specifically, we first straighten the *L* × *L* contact map matrices of all predicted structures into vectors and then apply principal component analysis (PCA) to obtain low-dimensional representations of the contact map. The number of principal components is selected to explain at least 90% of the total variance. We then apply K-medoids clustering on the PCA coordinates to extract *K* distinct structures. Note that AlphaFold2 is an ensemble model with five different sets of model weights, i.e., each MSA input generates five different structure predictions by AlphaFold2. Therefore, the selection of representative MSA subsets and all subsequent steps are performed independently for each of the five weight sets of AlphaFold2. Finally, for each of the *K* structures, we identify its corresponding MSA subset; these *K* MSA subsets will be used to extract the coevolutionary information.

#### Coevolutionary-Information-Aware MSA Sampling

By analyzing differences in coevolutionary information across representative MSA subsets, we develop an enhanced coevolution-aware sampling strategy to generate more diverse MSA subsets. The statistical model that we use for analyzing the coevolutionary information of the MSA is the Markov random field (MRF) model (*25*) with the parameter 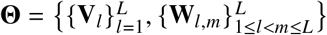,

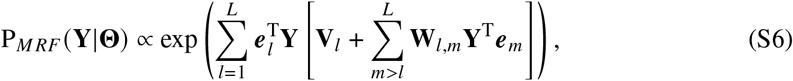

where **V**_*l*_ is a 22-dimensional vector capturing the marginal effect of the amino acid type at the *l*^*th*^ residue and **W**_*l, m*_ is a 22 × 22 symmetric matrix capturing the pairwise interaction effect between the *m*^*th*^ and *l*^*th*^ residues. **W**_*l, l*_ ≡ 0 for *l* = 1, …, *L*.

Sampling MSA subsets with diverse coevolutionary information can enable AlphaFold2 to more effectively explore the conformational space. To leverage this coevolutionary information effectively, we first estimate the MRF model for each of the representative MSA subsets extracted in the previous steps using GREMLIN (*25*). Suppose there are *K* MSA subsets, so that the MRF model is estimated *K* times. Then, for each pair of estimated MRF models, denoting their parameters by **Θ** and **Θ**^′^, we calculate the probability ratio P_*MRF*_ (**Y**_*i*_ |**Θ**)/P_*MRF*_ (**Y**_*i*_ |**Θ**^′^) across all sequences in the full MSA. A high ratio indicates that a sequence is more consistent with the coevolutionary information captured by **Θ** than **Θ**^′^. By pooling sequences with high probability ratios into a new MSA subset, we may strengthen their shared coevolutionary information. Finally, for each pair of estimated MRF models, we rank all sequences by their computed probability ratios and select the top *M* sequences, where *M* is a predetermined subset size. These *M* sequences are used as one input MSA subset for AlphaFold2 to generate new predictions. Since there are *K* (*K* − 1) such ordered pairs, this procedure generates *K* (*K* − 1) MSA subsets.

The steps of selecting representative MSA subsets and coevolutionary-information-aware MSA sampling are iterated in SMICE a few times (usually two or three rounds) to obtain the final structure predictions.

#### Identifying Regions of High Structural Variability

The conformational changes in fold-switching proteins can be broadly classified into two categories (*29*): (1) The structures of individual subregions remain largely unchanged, but their spatial arrangement shifts. This is often caused by the reorientation of a tail region (e.g., RelB protein). (2) The secondary or tertiary structure within a subregion changes.

Consequently, to locate regions of high structural variability from the predicted contact maps, we search for a contiguous region that meets the following criteria: it must exhibit high variance either in its intra-region contact distances or in its inter-region contact distances (i.e., its contacts with the rest of the protein), while the contact distances within the rest of the protein remain stable. Based on these criteria, we developed a stepwise algorithm to identify the variable region. The procedure is as follows:

(1) *Initialization*: The input is an ensemble of predicted contact maps for a protein of length *L*. We first compute an *L* × *L* matrix of variances, **∑**, where each entry **∑**_*ab*_ represents the variance of the contact distance between *a*^*th*^ and *b*^*th*^ residues across the ensemble. This matrix quantifies the positional variability in the structural predictions.
(2) *Identify Initial Candidate Region*: An initial candidate region is identified by scanning the protein sequence with a sliding window of a fixed window size, *w*_init_. For each sliding window, we compute the total pairwise variance of all residues *outside* the window and select the contiguous segment corresponding to the window that yields the minimum value. This ensures high variability of contacts, either within the segment or between the segment and the rest of the protein.
(3) *Calculating the Baseline Variability of Structural Conservation*: The baseline variability of structural conservation, denoted as *β*, is calculated as the average variance of all contact pairs where both residues lie outside the initial candidate region (i.e., pairs from outside the window).
(4) *Iterative Refinement*: The boundaries of the candidate region are refined through an iterative process that removes one residue at a time from either end of the sequence. In each iteration, we evaluate the consequence of removing the leftmost or the rightmost residue. The algorithm selects for removal the residue (leftmost or rightmost) that, when excluded, minimizes the total pairwise variance for all residues outside the new region.
(5) *Termination*: The iterative deletion process terminates when the mean variance of all contact pairs between the last-removed residue and the residues outside the current region exceeds 4*β*, or when the size of the candidate variable region reaches a predefined minimum. The detailed algorithm pseudocode is provided in Algorithm S3.

#### Representative Structures Extraction

We first filter out the low-quality predictions from SMICE’s sampling step based on a pLDDT threshold of 0.5 (recall that pLDDT is AlphaFold2’s built-in confidence measure). Next, to cluster the remaining predicted structures and identify the representative structures, we employed a stepwise clustering strategy based on structural similarity via TMscore. Since large-scale pairwise structural comparison based on TMscore is computationally prohibitive, we use Foldseek (*30*) to rapidly approximate TMscores. The procedure for stepwise clustering is as follows:

(1) The procedure begins with the highest-confidence structure (i.e., the one with the highest pLDDT score) among the five AlphaFold2 predictions generated from the full MSA. This structure serves as the initial “seed” for the first cluster.
(2) We calculate the TMscores between this seed structure and all structure predictions that were retained after filtering; all the TMscores are based on the specific region that includes the identified variable region and its left and right neighboring segments, each of which is taken to be half the length of the variable region. All structures with this TMscore greater than 0.85 relative to the seed structure are aggregated to form the first cluster. The structure with the highest pLDDT within this cluster is selected as its representative.
(3) To form each new cluster, the new cluster seed is chosen as the unclustered structure that has the lowest TMscore to the representative(s) of the previous cluster(s). Then, all unclustered structures with a TMscore > 0.85 to this new seed are grouped into a new cluster, and the structure with the highest pLDDT within the new cluster is taken as its representative.
(4) Step (3) is repeated until all structures are clustered.
(5) Finally, to ensure statistical reliability, any clusters containing fewer than three members are discarded as outliers.

This approach efficiently partitions the predicted structural ensemble into well-populated, high-confidence conformational states.

### Experiment Implementation Details

#### Setting Hyperparameters *τ* and Π

As described in the Method section, to promote diversity among the sampled MSA subsets, we vary the choices of **Π** in the prior distribution of the expected amino acid proportions.

We set these values in a data-driven manner. Specifically, for each sequence **Y**_*i*_ ∈ {**Y**_1_, …, **Y**_*n*_}, let 𝒮 (*i*) represent the indices of **Y**_*i*_’s N-nearest sequences among the full MSA using the Hamming distance. We compute the average proportions of amino acids in all residues across **Y**_*i*_’s neighbors, 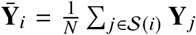. We then apply K-medoids clustering to these average amino acid proportions 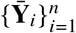 and obtain the resulting cluster centers. These cluster centers are used as *K* different choices for **Π**. In our experiment, we set *K* = 10. For MSAs with fewer than 100 sequences, we set *N* = 10, and for those with more than 100 sequences, we conduct the above procedure for both *N* = 10 and 30. Finally, we set the strength of the prior as *τ* = 0.5*N*.

#### Setting Hyperparameters in the Acceptance Probabilities

Since *λ* controls the homogeneity of the sequences within a sampled MSA subset, using multiple choices of *λ* could also lead to diverse MSA subsets in terms of conservation patterns in both the locations of conserved residues and the types of enriched amino acids. In particular, we choose *λ* from {0, 1, 2, 3}. Moreover, *C* in Eq.(S5) controls the overall acceptance probability. The common choice of *C* is set as 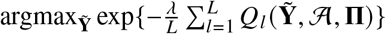 so that the acceptance probability can be no greater than 1.

#### Sequential Sampling Algorithm

The details of the sequential sampling algorithm are summarized in Algorithm S1. Notice that we set a maximum subset size *n*_max_ for efficient computation, and a minimum subset size *n*_min_ to ensure reliable AlphaFold2 predictions. The empirical suggestion of *n*_max_ is as follows: *n*_max_ = 20 if the full MSA size < 100, *n*_max_ = 100 if the full MSA size ∈ [100, 500], and *n*_max_ = 200 otherwise. We set the minimum subset size *n*_min_ = 10.

#### Estimating the MRF Model with GREMLIN

Given the MRF model in Eq.(S6) and sequences 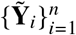, the log-likelihood of **Θ** is

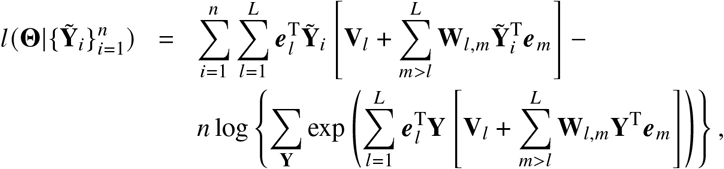

which is challenging to maximize directly due to the large number of values of **Y** in the summation. Given the high-dimensionality of **V**_*l*_ and **W**_*l, m*_, the GREMLIN method (*25*) considers the penalized

##### Algorithm S1

Sequential Sampling for MSA Subsets

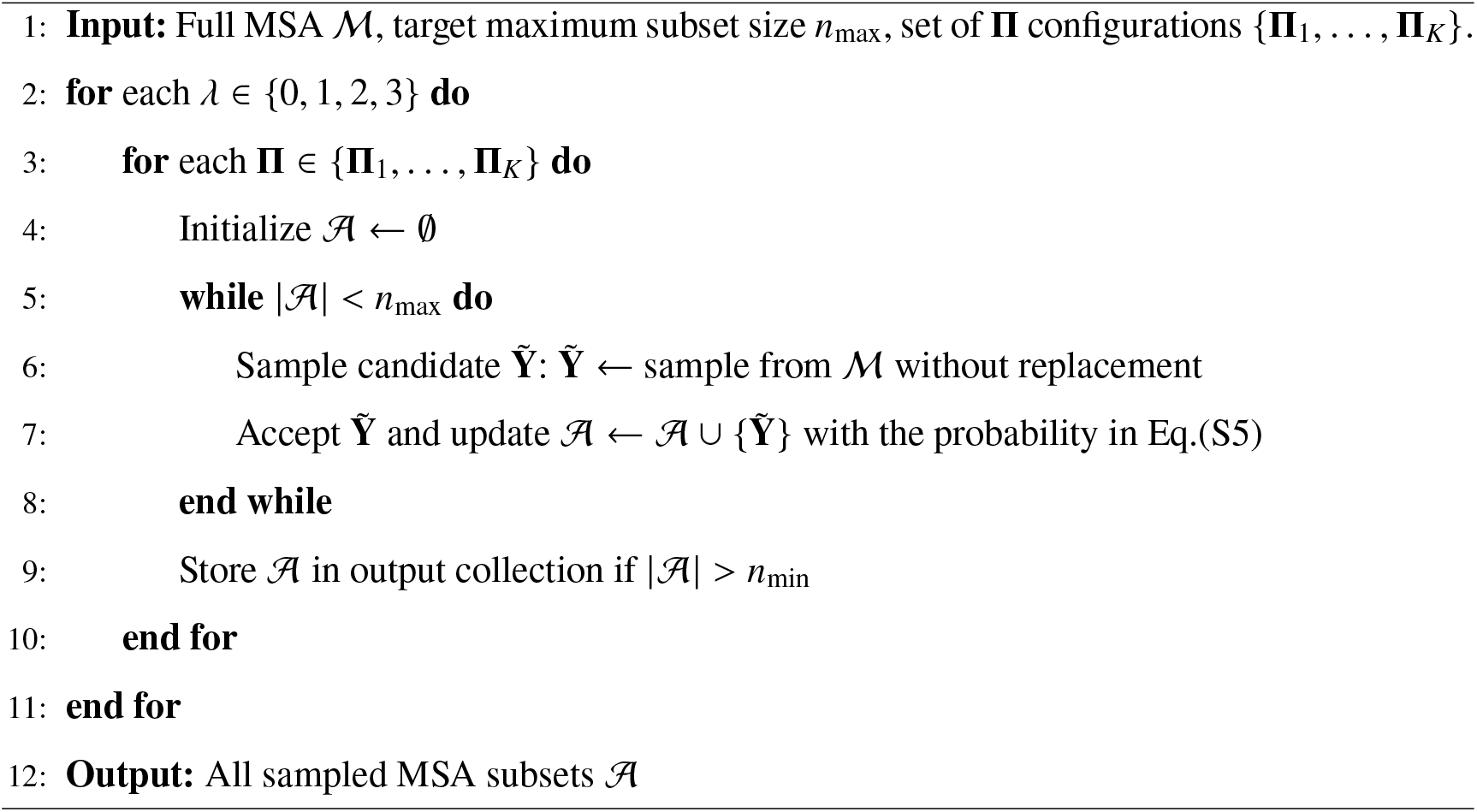

log pseudo-likelihood,

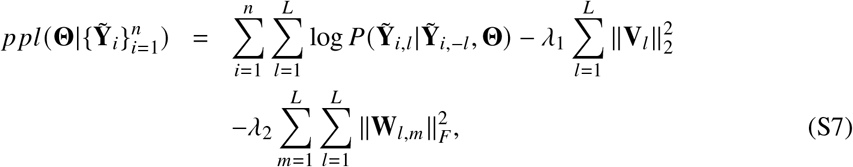

where 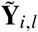 is the one-hot encoding of the amino acid at position *l* in the *i*^*th*^ sequence, and 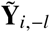 is the one-hot encoding of the amino acid at all other positions in the *i*^*th*^ sequence,

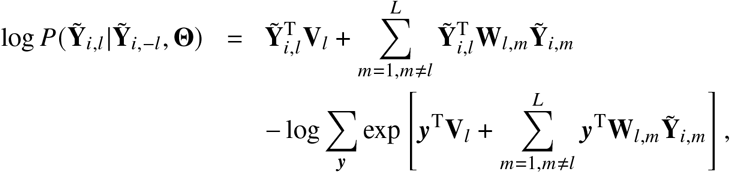

where ***y*** takes all one-hot encoding values of the amino acid types in the summation. The GREMLIN method then estimates **Θ** by maximizing Eq.(S7) with gradient descent using Adam (*31*); *λ*_1_ and *λ*_2_ are set to their default values in GREMLIN.

#### Coevolutionary-Information-Aware Sampling

The details of enhanced (coevolutionary-aware) sampling using coevolutionary information are summarized in Algorithm S2.

#### Identifying Regions of High Structural Variability

The details of identifying variable regions are summarized in Algorithm S3. To ensure the target variable region is entirely contained within the initial candidate region, we set an initial candidate region size that is sufficiently large. Additionally, the number of residues outside this region needs to be large enough to compute a reliable baseline of structural variance. Accordingly, we empirically set the initial candidate region size *w*_init_ = max(*L* − 40, 40), and the minimum variable region size *w*_min_ = 40, as stable protein domains are typically longer than 40 residues.

#### MSA Generation

MSAs were generated using MMseqs2 (*17*) implemented in ColabFold (*3*) by querying the UniRef30 database (*32*) of known sequences. Sequences were filtered to retain only those with ≥ 90% identity to the target sequence. The minimum coverage required for the MSA sequences is 75%. The MSA was enforced to have at most 4,096 sequences.

#### SMICE, AF-Cluster and Random Sampling

We consider 80 different hyperparameter configurations for MSA subset generation when the number of sequences exceeds 100, and 40 configurations when it is below 100. These configurations consist of four levels of *λ*, and 20 choices of **Π** for deep MSAs (sequence number ≥ 100) or 10 choices for shallow MSAs (sequence number < 100). For each configuration, we randomly draw four subsets using sequential sampling. We exclude any MSA subsets containing fewer than 10 sequences to ensure reliable AlphaFold2 predictions. Each remaining MSA subset is used to generate structures using ColabFold v1.5 with default settings, which outputs five predictions per run using AlphaFold2. No template structure is used in the prediction. The coevolutionary-information-aware sampling procedure is iteratively repeated twice (*n*_*iter*_ = 2), and we set the representative MSA subset *K* in Algorithm S2 as five, and the target MSA subset size as *M* = 20 if the full MSA has less than 300 sequences, and *M* = 100 if the full MSA has more than 300 sequences. This results in 200 predictions for deep MSA cases and 100 predictions for shallow MSA cases per iteration. In total, we obtain up to 1,800 predictions for deep MSAs and up to 900 predictions for shallow MSAs.

The results of *AF-Cluster* as well as *random sampling* are produced using the default setup of the AF-Cluster pipeline (*23*). In the pipeline, uniform random sampling without replacement is run up to 200 times on each MSA with sample sizes = 10 and 100 (if the full MSA has more than 100 sequences). The predictions are made using ColabFold v1.5 with default settings.

#### Computing comparative metrics

To ensure a fair comparison between methods when comparing the best prediction result for each method on each fold-switching protein with max-TMscore1, max-TMscore2, and minimax-TMscore, we standardized the sizes of prediction sets as follows:

For AF-Cluster, we used the full prediction set, given its number of predictions is inherently determined by the number of clusters.

For Random Sampling, we matched the prediction set size to that of AF-Cluster in each case.

For SMICE, if its prediction set was smaller than AF-Cluster’s, we used all available predictions. If it was larger, we repeatedly subsampled SMICE predictions 500 times to match the number of predictions of AF-Cluster, and calculate the averaged metrics across all repetitions.

### Additional Experiments

#### MSA Subsets with Different Sequence Diversity Predict Distinct Conformations

Using the thioesterase SP 1851 (PDB: 4zrb) (*33*) as an example, we investigated how the diversity of MSA subsets affects conformational predictions. SP 1851 forms a homotetramer, and each chain can adopt two distinct conformational states: 4zrbC and 4zrbH (Figure S1b). We computed the TMscores on the fsr and neighboring segments of the predicted structures generated by SMICE to both of the conformational states.

We found that the conformational prediction preference depends on the parameter *λ* of the sequential sampling step, which controls the sequence diversity of the MSA sampled subset. As shown in Figure S1a,c, MSA subsets that successfully predicted the 4zrbC conformation were predicted from MSA subsets with more diverse sequences (lower *λ*), whereas subsets that predicted the 4zrbH conformation consisted of less diverse sequences (higher *λ*). This observation demonstrates that sampling MSA subsets with multiple levels of sequence diversity ensures broader coverage of the conformational space.

#### Coevolutionary Information Enables SMICE’s Prediction of the Fold-switching State

Using the di-haem cytochrome c peroxidase (CCP) protein (*34*) as an example, we investigated how enhancing coevolutionary-information diversity across the MSA subsets helps the prediction of the alternative conformation. The CCP protein adopts two conformational states in the mixed-valence form (PDB: 2c1v, Figure S3a) and in the oxidized form (PDB: 2c1u, Figure S3b). In the fold-switching region (highlighted by the dashed circle in Figure S3), 2c1v forms a loop, whereas the corresponding region in 2c1u contains segments of *β*-sheet and *α*-helix. As both are homodimers, we evaluated predictions against chain B of 2c1v and chain C of 2c1u. As shown in Figure S2a, predictions from sequential sampling only, without enhanced coevolutionary-aware sampling, are unable to capture the fold-switching state 2c1uC, whereas predictions from the full SMICE sampling step successfully capture both conformation states. To analyze the underlying cause, we characterized the coevolutionary information for each MSA subset using the Markov random field model (Eq. S6), which estimates the residue-to-residue coupling matrix **W**. UMAP visualization of these **W** matrices (Figure S2b) revealed that the diversity of coevolutionary information among MSA subsets generated by the full SMICE sampling is substantially greater than among those from sequential sampling only. Moreover, the **W** matrices of MSA subsets that predicted the fold-switching state form a distinct, separate cluster as highlighted in the dashed circle in Figure S2b. This increased diversity in coevolutionary information appears to be the key that enables the prediction of the fold-switching state. For completeness, Figure S3 compares the SMICE’s top predictions with those from AF-Cluster and random sampling, neither of which predicts the fold-switching state 2c1uC.

#### Ablation Study of SMICE’s sampling step

We estimate the importance of key components of SMICE’s sampling step by considering ablations on SMICE:

##### SMICE (full)

SMICE (full), as described in the paper, returns the full set of predictions generated by SMICE’s sampling step, including those from the sequential sampling and the coevolutionary-information-aware MSA sampling.

##### SMICE (sequential only)

SMICE (sequential only) returns the predictions from sequential sampling only, *without* conducting enhanced coevolutionary-information-aware MSA sampling.

##### SMICE (no coevol)

For SMICE (no coevol), we remove the coevolutionary term in the MRF model when conducting the enhanced sampling, i.e., we replace Eq. (S6) with

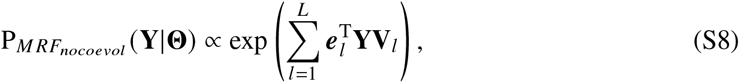

and estimate **V**_*l*_ with the corresponding penalized log pseudo-likelihood,

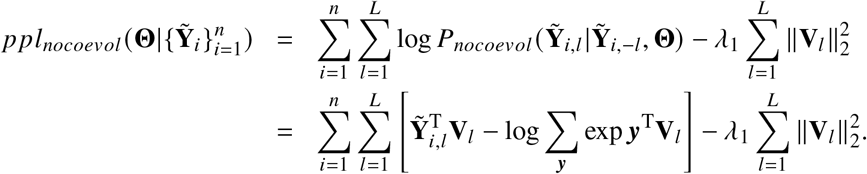

In other words, SMICE (no coevol) returns the combined predictions from sequential sampling and enhanced sampling using the MRF model but without the coevolutionary term.

We performed bootstrap hypothesis testing to compare SMICE (full) against the two ablations SMICE (sequential only) and SMICE (no coevol) on three metrics: max-TMscore1, max-TMscore2, and minimax-TMscore. The TMscore is calculated on the fsr plus its neighboring segments. To ensure fair comparison, when SMICE (sequential only) generated fewer predictions, we repeatedly (500 times) randomly subsampled SMICE (full) predictions to match the sample size of SMICE (sequential only), then averaged the metrics across these subsamples. For each protein, we computed the difference in each metric (e.g., max-TMscore1(full) - max-TMscore1(sequential only)). We tested the null hypothesis that the mean difference ≤ 0, where a p-value < 0.05 would indicate that SMICE (full) performs significantly better than the ablation on that metric. We performed 10,000 bootstrap resamples of the fold-switching cases to estimate the p-values as reported in Table S1 and Table S2. The statistical analysis reveals that SMICE (full) significantly improves all three metrics over SMICE (sequential only). For the coevolutionary information ablation comparison, SMICE (full) does not show significant improvement over SMICE (no coevol) in predicting the ground state. However, it demonstrates significant improvement for predicting the (more challenging to predict) fold-switching state (max-TMscore2 p-value = 0.036).

### Confidence Metrics Failed to Rank the Predictions of Fold-switching Proteins

AlphaFold2’s built-in confidence metrics (pLDDT, predicted TMscore (PTM), and predicted aligned error (PAE)) are widely used for ranking predictions. While higher pLDDT/PTM (with range 0-1) and lower PAE (with range ≥ 0) typically indicate higher confidence, we find these scores unreliable for directly ranking predictions of fold-switching proteins.

Our analysis of SMICE’s sampling step for the benchmark set of fold-switching proteins shows a general trend: pLDDT and PTM positively correlate with prediction accuracy (measured by TMscore1, TMscore2, and max-TMscore (where TMscore is calculated on the fsr plus its neighboring segments), and PAE shows a negative relationship (Fig. S4). However, pLDDT and PTM are often overestimated, and PAE is usually underestimated. For example, predictions with high pLDDT scores (close to 1.0) exhibit a wide range of max-TMscore values (0.4–1.0).

This issue becomes clear when examining individual fold-switching proteins. For example, a fragment of the parainfluenza virus 5 fusion protein has two conformations (Fold1: 1svfC, Fold2: 4wsgC), and the Spearman correlation between the predictions’ pLDDT and max-TMscore is statistically significantly negative (correlation value = -0.228, p-value = 1.146 × 10^−15^; Fig. S5a). This counterintuitive result extends to a substantial subset of all fold-switching proteins for other confidence metrics (PTM, PAE) across all TMscore measures (TMscore1, TMscore2, and max-TMscore) (Fig. S5b).

We attribute this phenomenon to a systematic bias between the confidence scores of the two conformations. In the example above, predictions resembling 4wsgC (red) have systematically higher pLDDT but lower max-TMscore than those resembling 1svfC (blue). Consequently, while the relationship between pLDDT and max-TMscore is positive within the group of predictions similar to 4wsgC, the combination of both groups creates an overall negative relationship.

Nevertheless, pLDDT remains a useful metric for filtering out low-quality predictions, as low pLDDT values consistently correspond to low max-TMscores (Fig. S4a).

#### Selecting the pLDDT Threshold for Prediction Filtering

We compared the full prediction set from the SMICE’s sampling step against the prediction sets filtered by pLDDT thresholds (0.3, 0.4, 0.5, 0.6) to select the best threshold based on six metrics from our benchmark study. The three metrics, including max-TMscore1, max-TMscore2, and minimax-TMscore, assess the accuracy of the top prediction for each prediction set. Since filtering removes predictions, the values for these metrics in the filtered prediction set cannot exceed those of the full set. Therefore, an optimal threshold is one that preserves these scores as close as possible to their values in the full prediction set. The other three metrics, including the proportion for predicting Fold1, the proportion for predicting Fold2, and the minimum proportion for both folds, measure the overall accuracy of the prediction set. For these metrics, the goal is to choose a threshold that maximizes their values.

Given this, we found that pLDDT thresholds of 0.3, 0.4, and 0.5 performed almost equally well in the top predictions’ accuracy as the full prediction set (Fig. S6a-c). Furthermore, a threshold of 0.5 provided the greatest improvement in overall prediction accuracy, making it our chosen value (Fig. S6d-f). By uniformly applying a pLDDT threshold of 0.5 to all predictions, we successfully filtered out approximately 24% of the predictions.

In contrast, filtering based on either PTM or PAE did not yield a significant improvement in overall prediction accuracy, as shown in Fig. S7.

##### Algorithm S2

Enhanced MSA Sampling using Coevolutionary Information

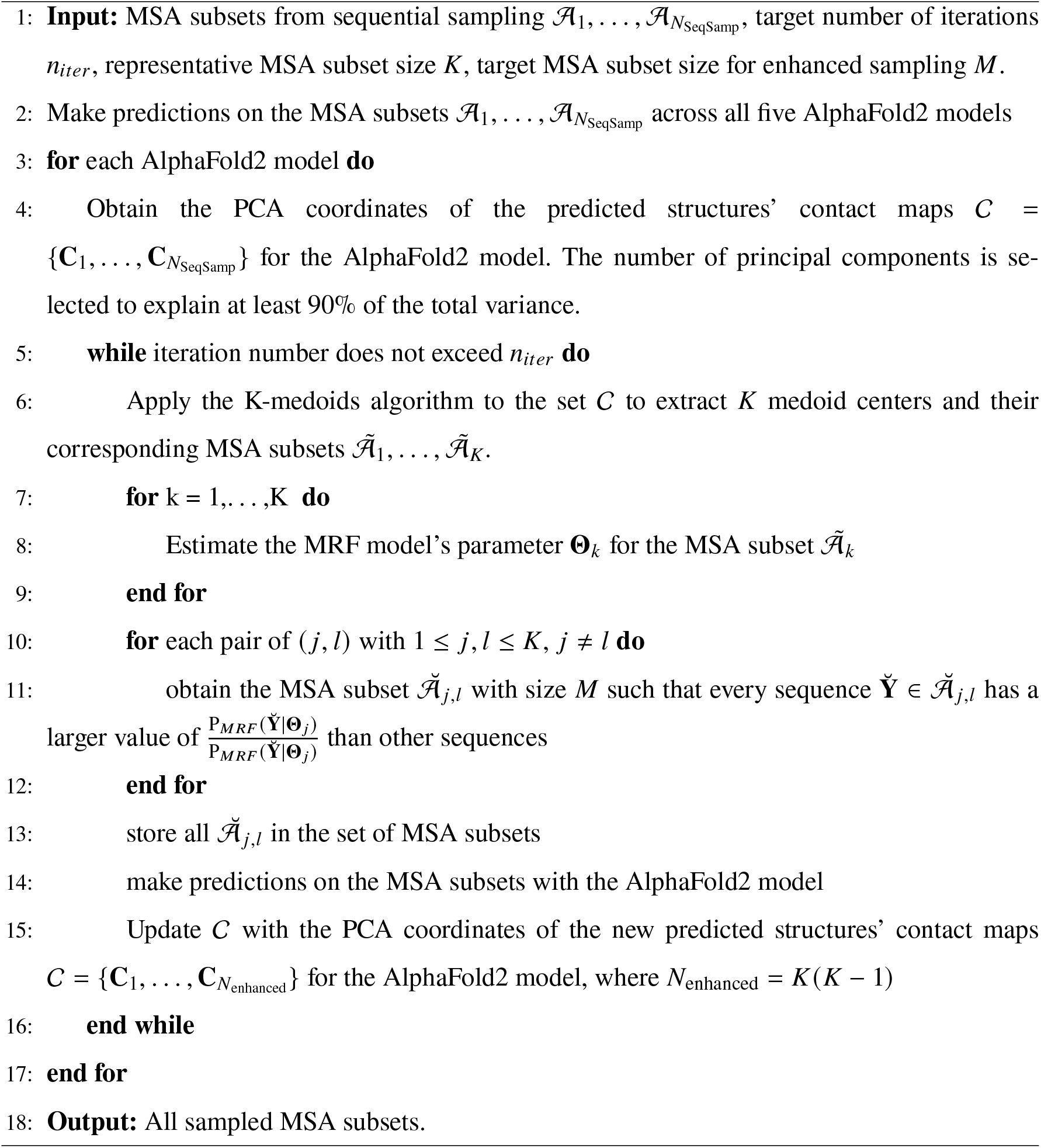

##### Algorithm S3

Identifying Region of High Structural Variability

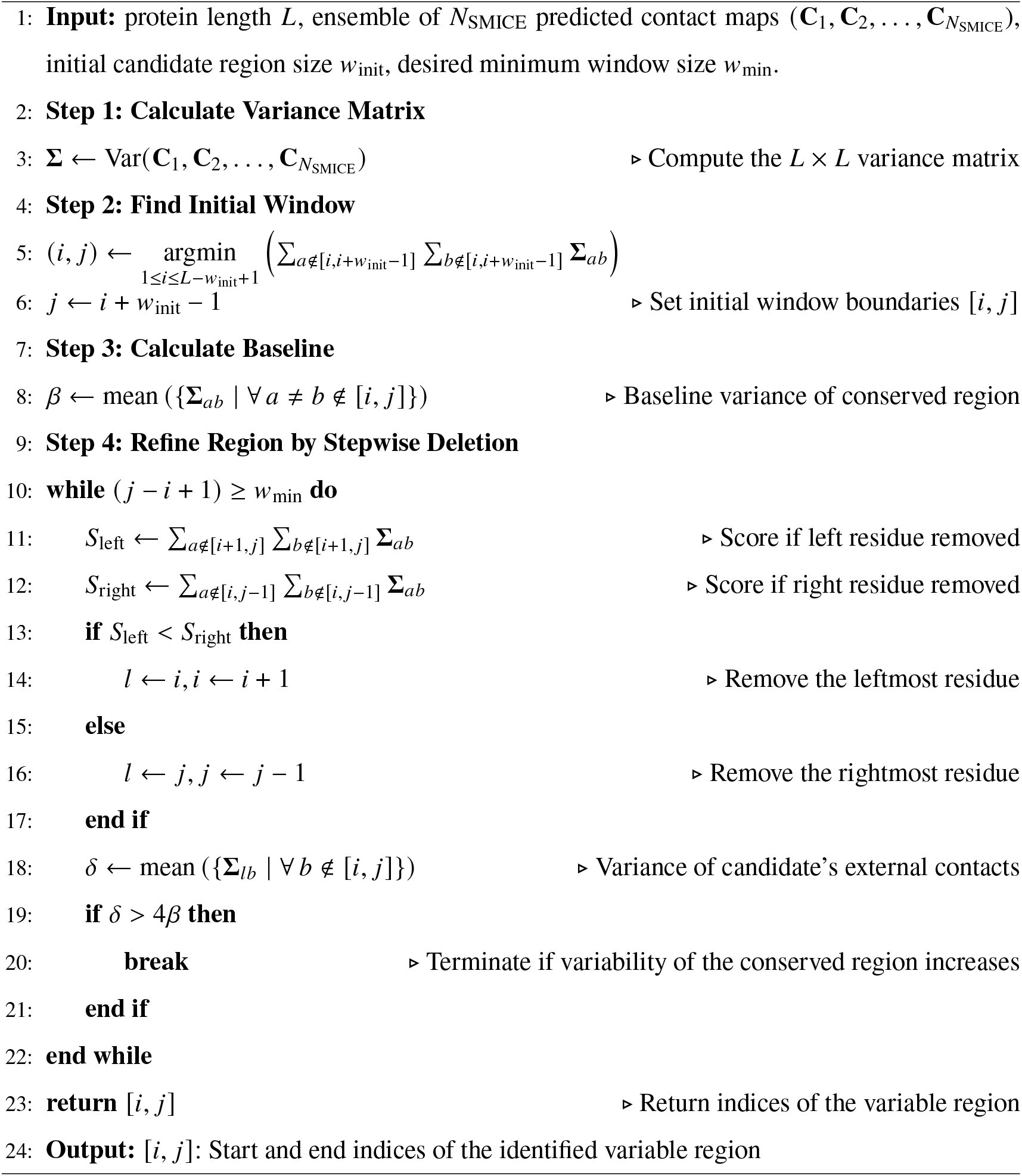

---

**Figure S1.**
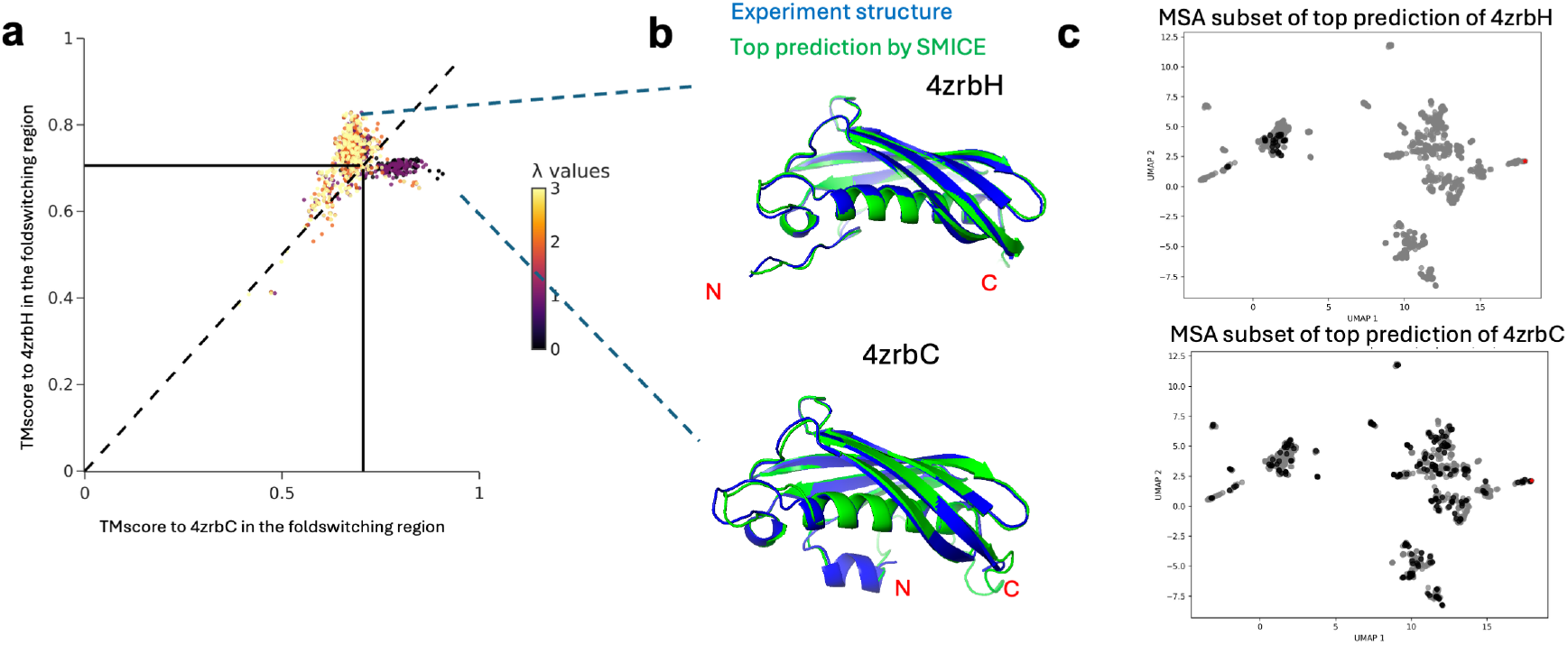
**a**, TMscores on the fold-switching region plus the neighboring segments of predictions generated by SMICE’s sampling step. Black vertical and horizontal lines indicate TMscores between SP 1851’s two experimentally determined conformations. Predictions are colored according to the values of the sequence diversity parameter *λ*, which is used by SMICE during the generation of their respective MSA subsets. *λ* explicitly controls the degree of sequence diversity in the sampled MSA subsets, with lower values selecting more diverse sequences. **b**, Crystal structures of SP 1851 in its ground (PDB: 4zrbC) and fold-switching (PDB: 4zrbH) states alongside their top SMICE-predicted structures. **c**, UMAP visualization of the MSA sequences and subsets that make the top predictions of the two conformational states. The sequences sampled in the MSA subset are colored in black, and the unsampled sequences are colored in gray.

**Figure S2.**
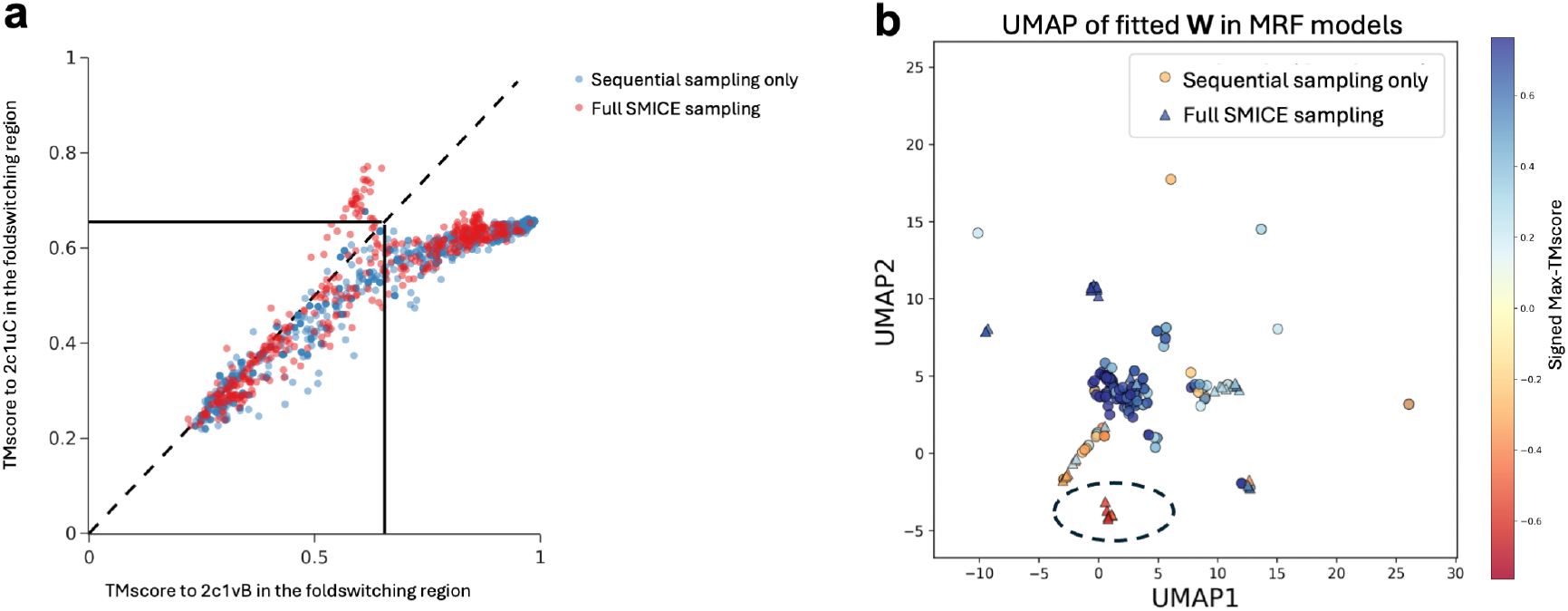
**a**, Comparison between predictions on CCP’s conformational states (PDB: 2c1vB and 2c1uC) from sequential sampling only (blue) and the full SMICE sampling (red). Black vertical and horizontal lines indicate TMscores between the two conformational states. **b**, UMAP visualization of the distribution of coevolutionary information, measured by **W** in the Markov random field model Eq.(S6), from sequential sampling only and the full SMICE sampling.

**Figure S3.**
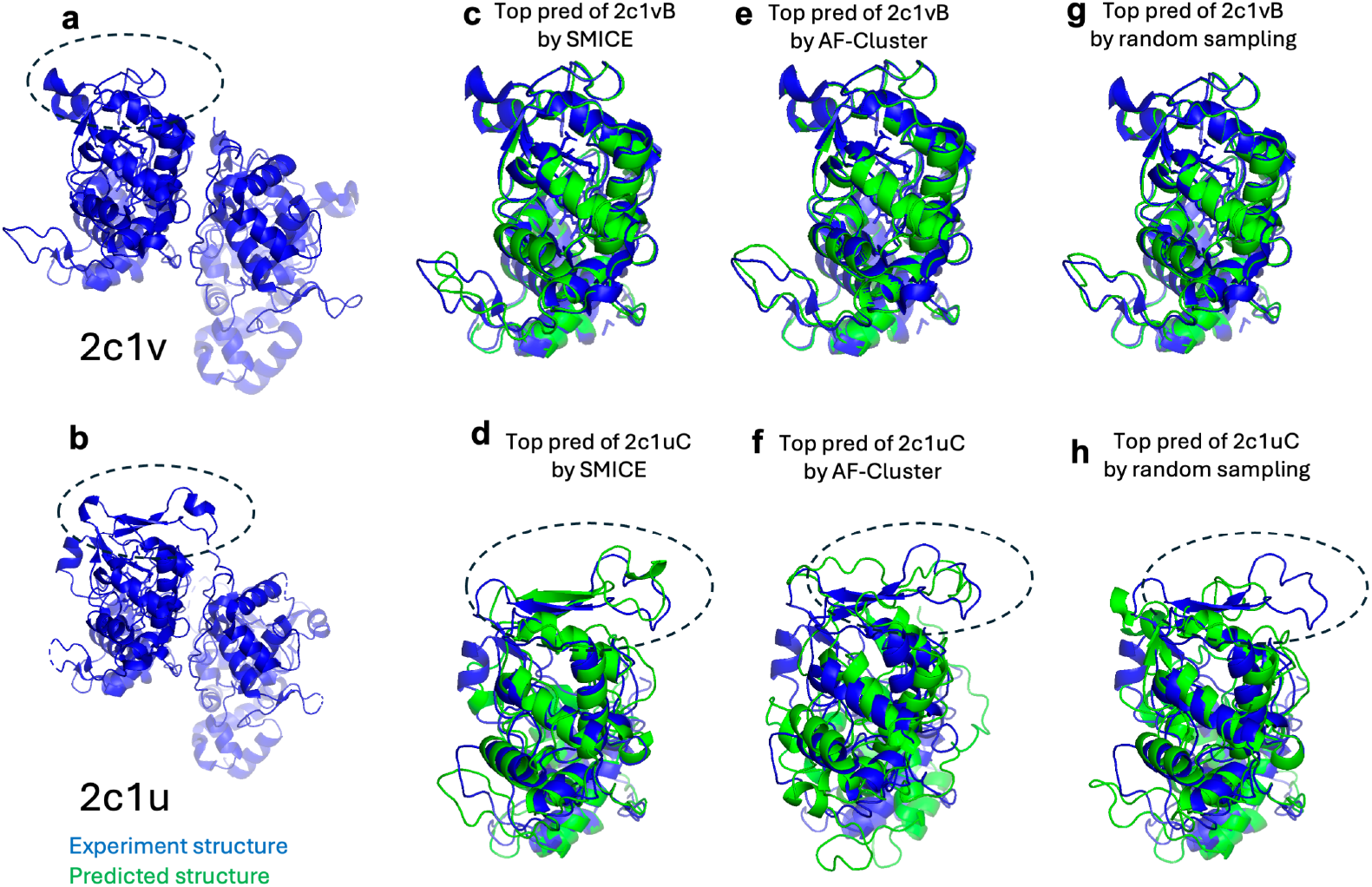
SMICE samples both known conformations of di-haem cytochrome c peroxidase (CCP). Crystal structures of CCPs in the mixed-valence form (PDB: 2c1v) (**a**) and the oxidized form (PDB: 2c1u) (**b**). **c-d**, top-ranked predictions of 2c1vB and 2c1uC by SMICE, respectively. **e-f**, top-ranked predictions of 2c1vB and 2c1uC by AF-Cluster, respectively. **g-h**, top-ranked predictions of 2c1vB and 2c1uC by random sampling, respectively.

**Figure S4.**
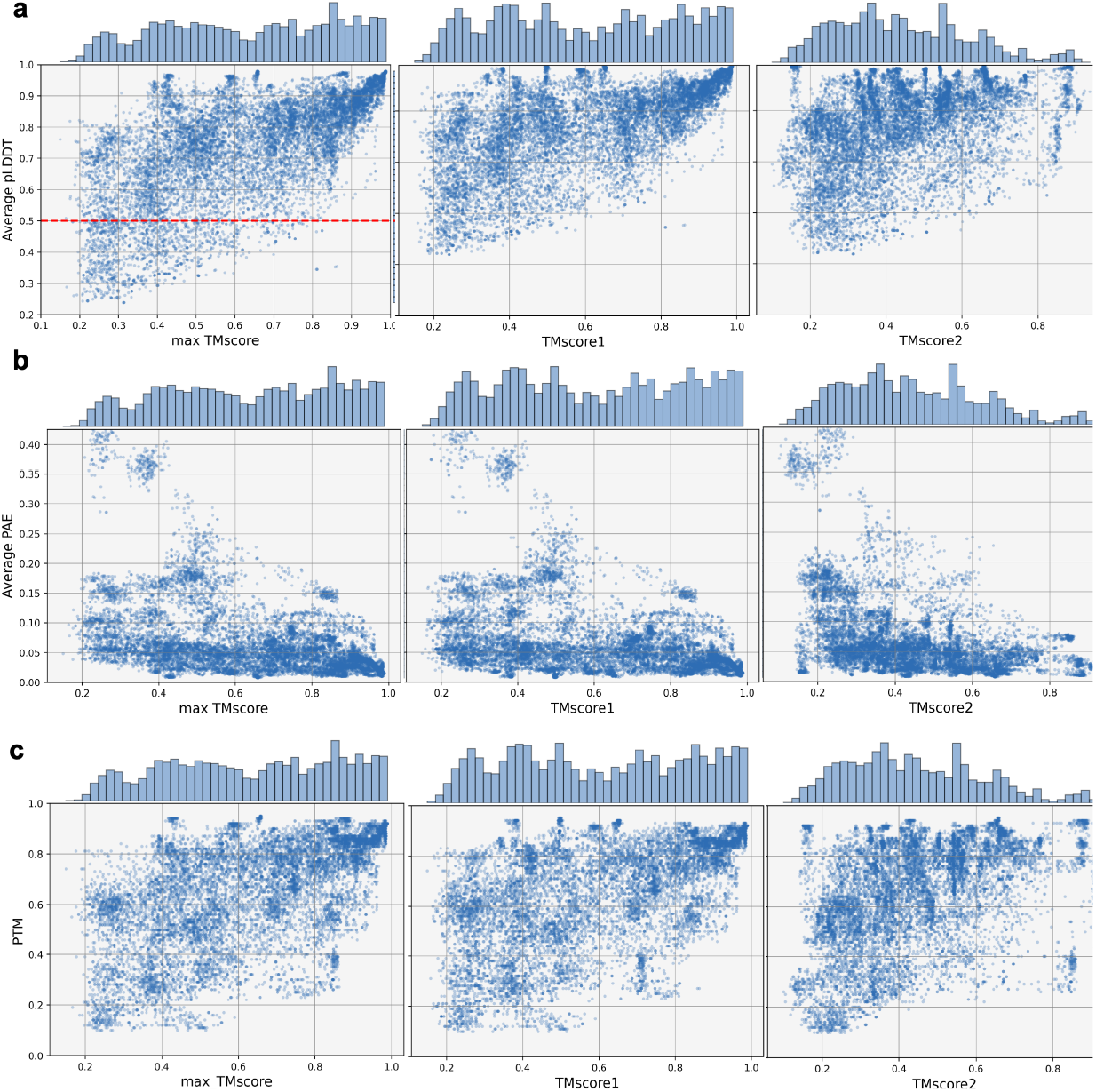
**a**, pLDDT versus max-TMscore, TMscore1, and TMscore2 for all fold-switching protein predictions. The filtering threshold (pLDDT = 0.5) is visualized with the red dashed line. **b**, PAE versus max-TMscore, TMscore1, and TMscore2 for all fold-switching protein predictions. **c**, PTM versus max-TMscore, TMscore1, and TMscore2 for all fold-switching protein predictions.

**Figure S5.**
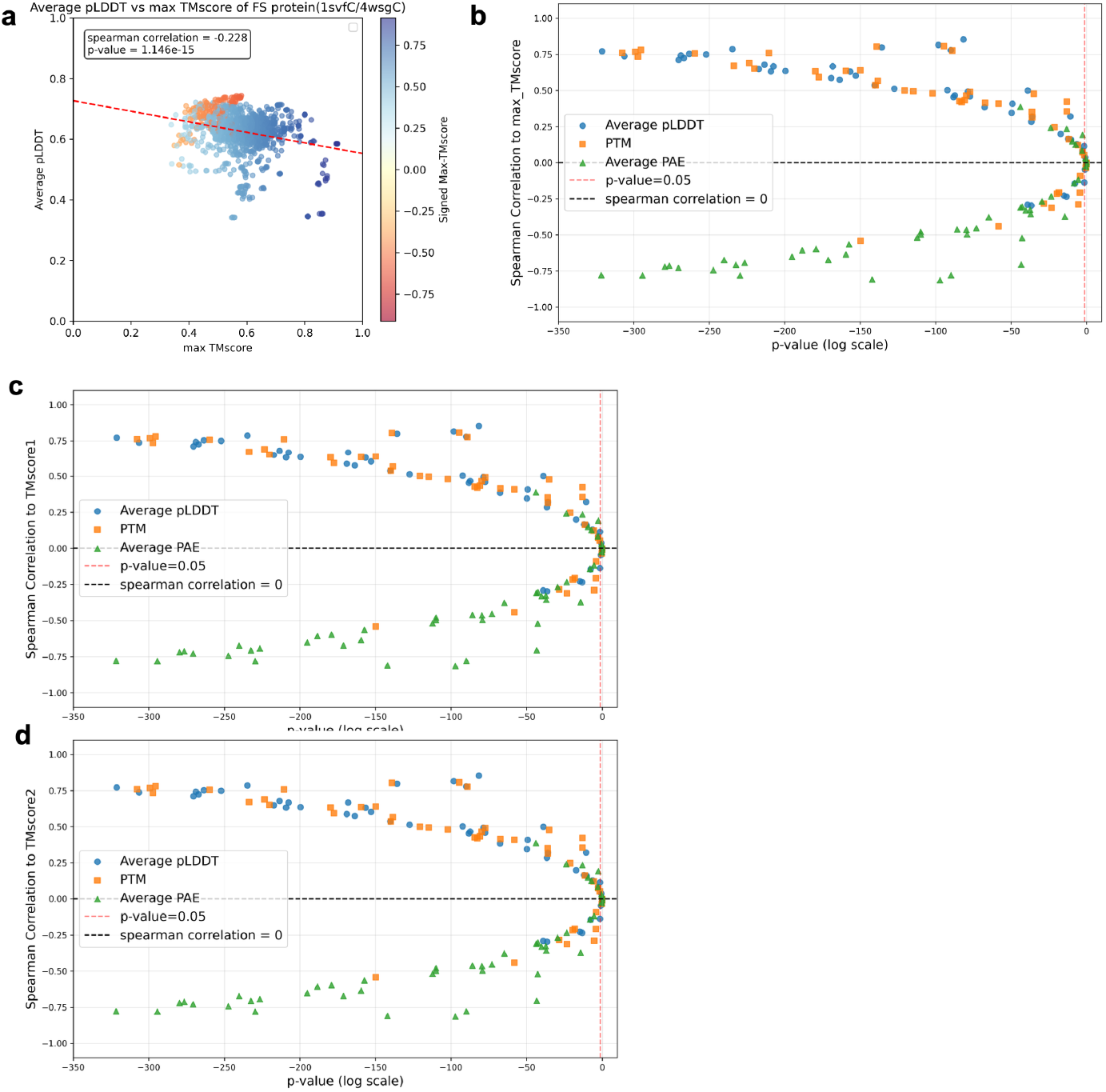
**a**, For the fold-switching protein with the two conformations (PDB IDs: 1svfC and 4wsgC), the pLDDT scores and max-TMscore exhibit a negative correlation (red dashed line represents the regression fit of the pLDDT scores against max-TMscore). Predictions are colored by the signed max-TMscore (sign[TMscore1 – TMscore2] × max-TMscore). **b**, Spearman correlation between confidence metrics (pLDDT, PAE, and PTM) and max-TMscore versus the correlation test p-values across all fold-switching proteins. The red dashed line marks significance (p-value = 0.05). **c**, Spearman correlation between confidence metrics (pLDDT, PAE, and PTM) and TMscore1 versus the correlation test p-values across all fold-switching proteins. The red dashed line marks significance (p-value = 0.05). **d**, Spearman correlation between confidence metrics (pLDDT, PAE, and PTM) and TMscore2 versus the correlation test p-values across all fold-switching proteins. The red dashed line marks significance (p-value = 0.05).

**Figure S6.**
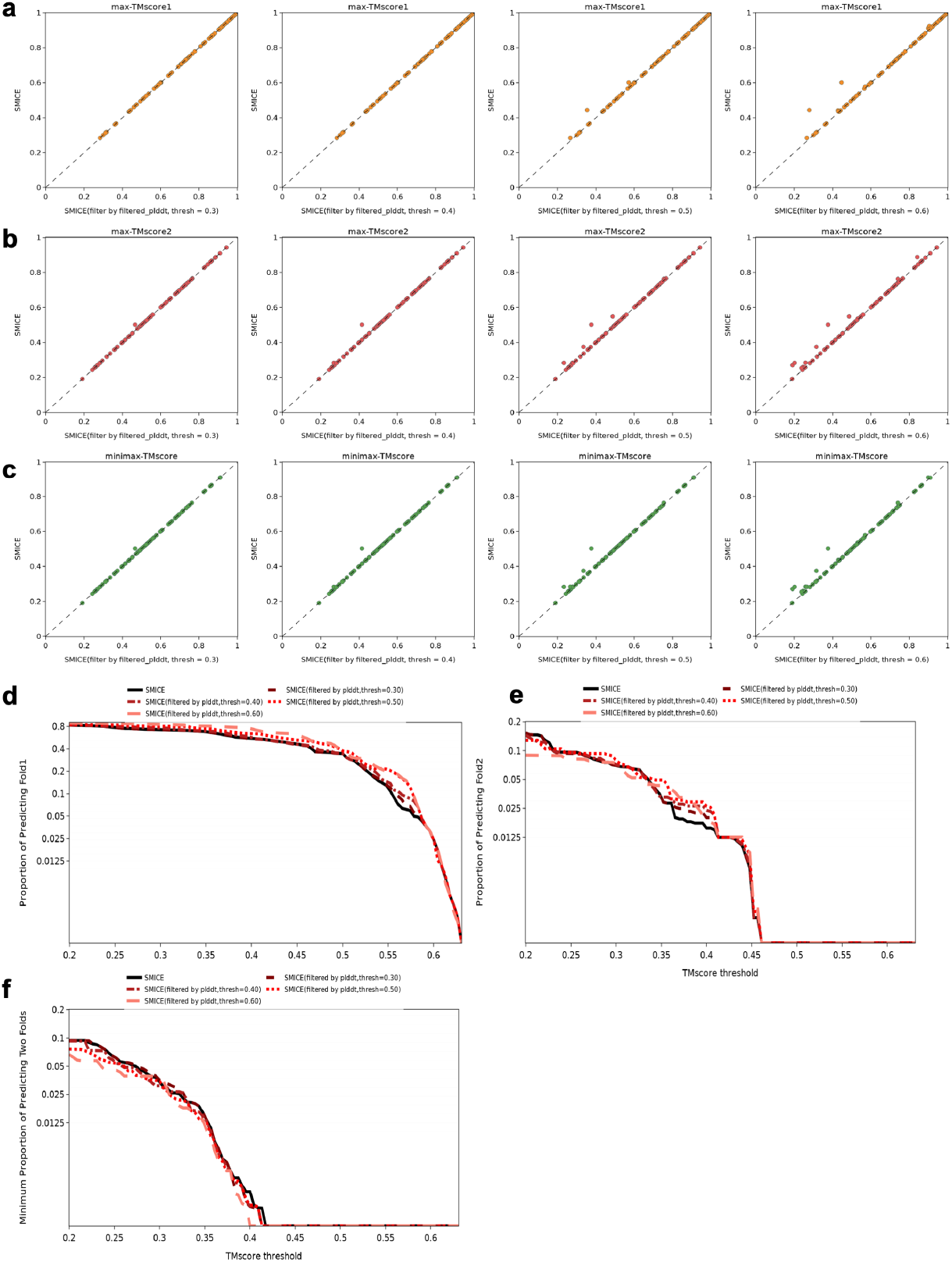
Filtering prediction sets from the SMICE’s sampling step based on pLDDT thresholds (greater than 0.3, 0.4, 0.5, 0.6). **a-c**, Comparison of the full prediction set against the filtered prediction set on the max-TMscore1, max-TMscore2, and minimax-TMscore. **d-f**, Comparison of the full prediction set against the filtered prediction set on the proportions of predicting Fold1, the proportions of predicting Fold2, and the minimum proportions of predicting both folds.

**Figure S7.**
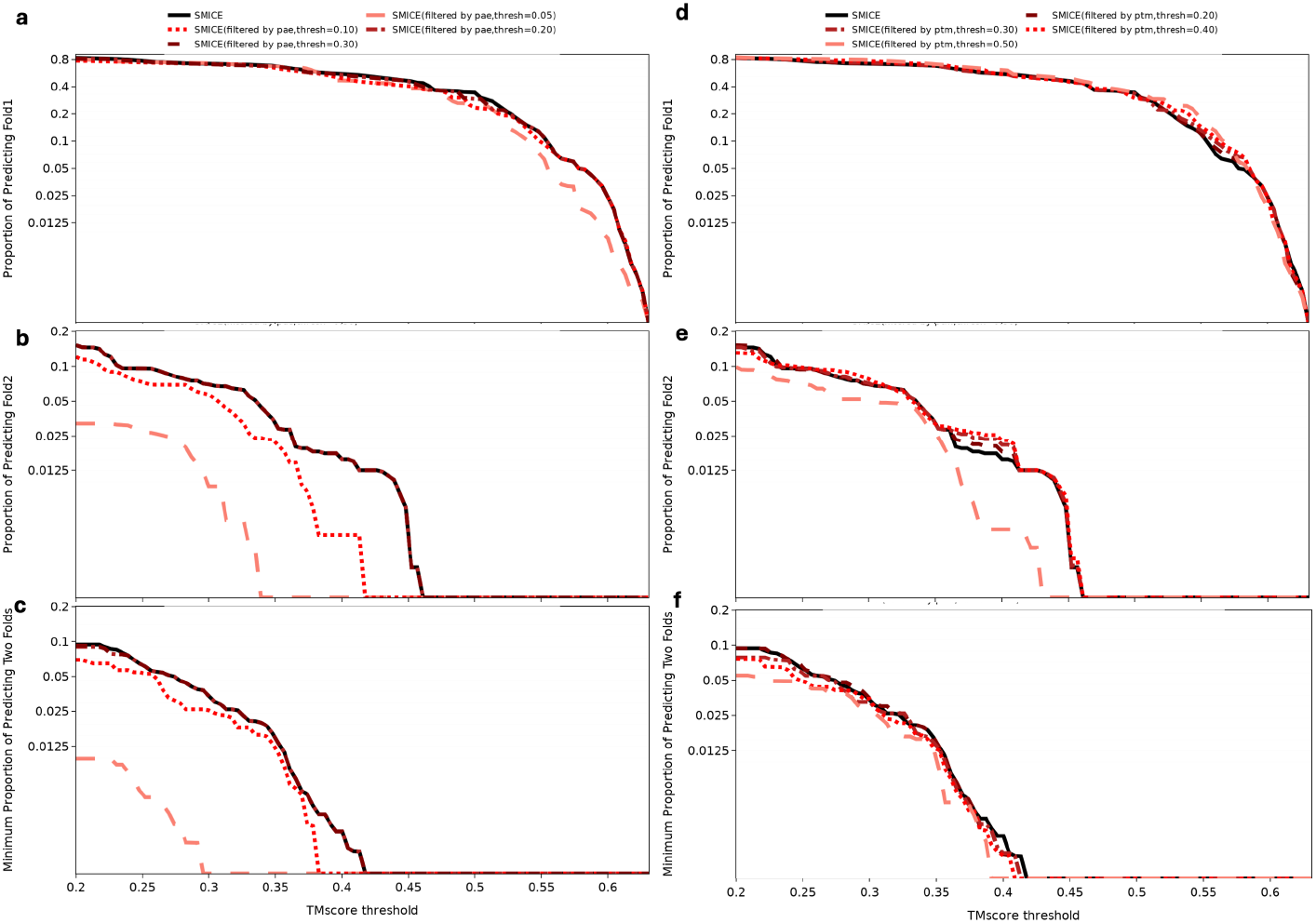
Filtering prediction sets from the SMICE’s sampling step based on PAE and PTM thresholds. **a-c**, Comparison of the full prediction set against sets filtered by PAE thresholds (< 0.05, 0.1, 0.2, 0.3), showing the proportions of predicting Fold1, the proportions of predicting Fold2, and the minimum proportions of predicting both folds. **d-f**. Comparison of the full prediction set against sets filtered by PTM thresholds (> 0.2, 0.3, 0.4, 0.5), showing the proportions of predicting Fold1, the proportions of predicting Fold2, and the minimum proportions of predicting both folds.

**Figure S8.**
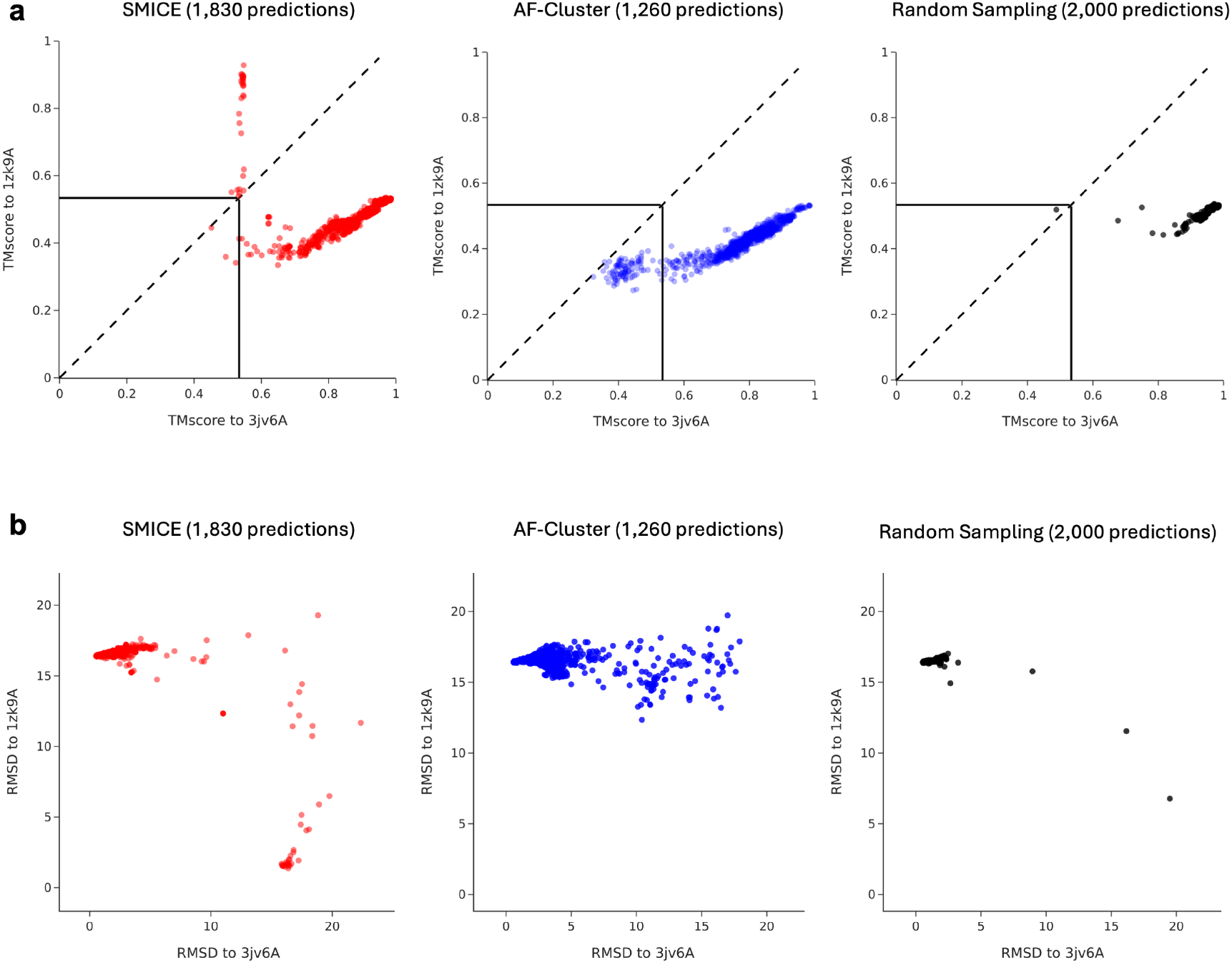
Results of SMICE’s sampling step on the fold-switching protein RelB. **a**, TMscores of the whole protein structure for the AlphaFold2 predictions generated by the SMICE sampling step (red), AF-Cluster (blue), and random sampling (black). Black vertical and horizontal lines indicate the whole-structure TMscores between RelB’s two experimentally determined conformations, 3jv6A and 1zk9A. **b**, RMSDs of the whole protein structure for AlphaFold2 predictions generated by SMICE sampling (red), AF-Cluster (blue), and random sampling (black).

**Figure S9.**
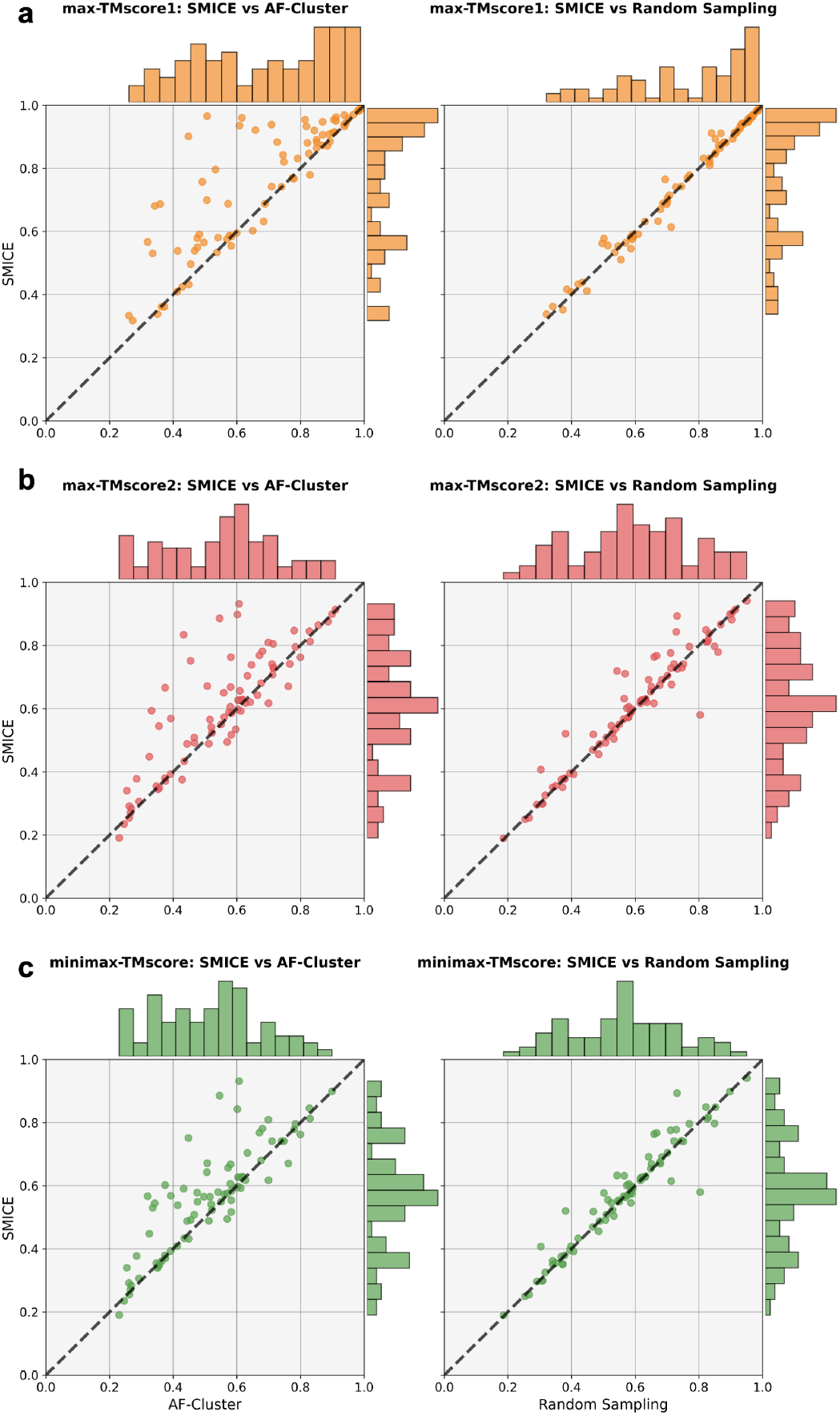
Comparison of the max-TMscore1, max-TMscore2, and minimax-TMscore for the full set of predictions generated by SMICE’s sampling step against AF-Cluster and Random Sampling. Each point represents one fold-switching protein.

**Table S1.**
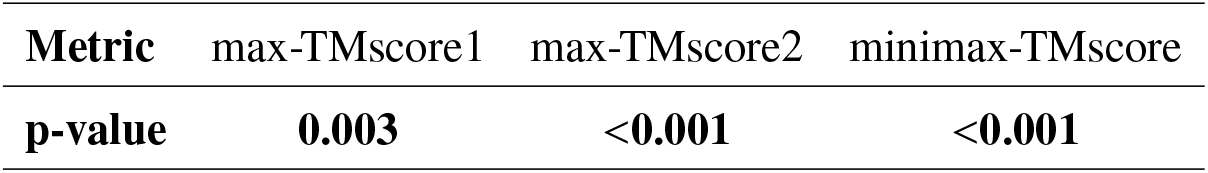
Statistical analysis comparing SMICE (full) versus SMICE (sequential only). P-values from bootstrap test (null hypothesis: mean of a metric of SMICE (full) ≤ mean of the corresponding metric of SMICE (sequential only)).

| **Metric** | max-TMscore1 | max-TMscore2 | minimax-TMscore |
| --- | --- | --- | --- |
| **p-value** | **0.003** | **<0.001** | **<0.001** |

**Table S2.** Statistical analysis comparing SMICE (full) versus SMICE (no coevol). P-values from bootstrap test (null hypothesis: mean of a metric of SMICE (full) ≤ mean of the corresponding metric of SMICE (no coevol)).

| **Metric** | max-TMscore1 | max-TMscore2 | minimax-TMscore |
| --- | --- | --- | --- |
| **p-value** | **0.746** | **0.036** | **0.035** |

